# Mid-zone hepatocytes trade proliferation for survival via Atf4-Chop axis in early acute liver injury

**DOI:** 10.1101/2025.08.21.671501

**Authors:** Yaying Zhu, Chengxiang Deng, Bo Chen, Jia He, Yanan Liu, Shan Lei, Weiju Lu, Cheng Peng, Zhao Shan

**Affiliations:** Yunnan Key Laboratory of Cell Metabolism and Diseases, Center for Life Sciences, School of Life Sciences, Yunnan University, Kunming, China, 650500

**Keywords:** Hepatocytes, Atf4-Chop axis, Btg2, acetaminophen-induced liver injury

## Abstract

Hepatocytes undergo extensive proliferation to facilitate liver repair after injury, yet early adaptive changes prior to proliferation remain unclear. Here, we report that during early acetaminophen (APAP)-induced liver injury, hepatocytes exhibit transient proliferation suppression that is most pronounced in mid-zone hepatocytes, consistent with zonal APAP metabolism. While spatial transcriptomics (ST) provided robust evidence for this arrest in the mid-zone, support for a similar arrest in pericentral hepatocytes was limited. Integrating ST with immunohistochemistry and functional studies, we identified a unique mid-zone stress-response program centered on the Atf4–Chop axis, which suppresses proliferation via the cell cycle inhibitor Btg2. Together, our findings support a model in which mid-zone hepatocytes transiently prioritize stress adaptation over proliferation, thereby preserving regenerative capacity for subsequent liver repair.

## Introduction

Acetaminophen (APAP) overdose is the leading cause of drug-induced acute liver failure worldwide, resulting in over 10,000 hospitalizations and approximately 500 deaths annually in the United States (Ostapowicz, Fontana et al. 2002, Larson, Polson et al. 2005, Lee 2007, Bernal and Wendon 2013). The hepatotoxicity of APAP is initiated by its metabolic activation through cytochrome P450 enzymes, primarily in pericentral hepatocytes, generating the reactive metabolite N-acetyl-p-benzoquinone imine (NAPQI) (Dahlin, Miwa et al. 1984, Hinson, Reid et al. 2004, Jack A. Hinson 2010, Tujios and Fontana 2011). NAPQI depletes glutathione and forms protein adducts, leading to oxidative stress and mitochondrial dysfunction that culminate in centrilobular necrosis (Dahlin, Miwa et al. 1984, Hinson, Reid et al. 2004, Jack A. Hinson 2010, Tujios and Fontana 2011). Although the mechanisms of APAP bioactivation and toxicity are well-characterized, the early adaptive responses of hepatocytes—particularly their proliferative dynamics across lobular zones—remain poorly understood.

The liver is organized into functional hexagonal units termed lobules, where hepatocytes display spatial heterogeneity across three distinct zones: periportal (PP, zone 1), mid-zonal (Mid, zone 2), and pericentral (PC, zone 3) (Ben-Moshe, Veg et al. 2022, Wang, Wang et al. 2024, Wu, Shentu et al. 2024). These zones exhibit unique gene expression profiles and metabolic specializations(Halpern, Shenhav et al. 2017). Pericentral hepatocytes (zone 3), expressing high levels of cytochrome P450 enzymes, are the primary site of APAP-induced necrosis (Wang, Wang et al. 2024). However, recent spatial profiling, functional, and lineage tracing studies have identified midzonal hepatocytes (zone 2) as the predominant source of new hepatocytes during both homeostasis and repair, demonstrating significant proliferative capacity and metabolic plasticity (Chembazhi, Bangru et al. 2021, He, Pu et al. 2021, Wei, Wang et al. 2021). Despite these advances, how mid-zone hepatocytes dynamically adapt during the initial phase of acute stress remains unknown.

A key regulator of cellular stress adaptation is the integrated stress response (ISR), which phosphorylates eukaryotic initiation factor 2α (eIF2α) to suppress global translation while selectively upregulating stress-responsive genes, including activating transcription factor 4 (Atf4) and its downstream target pro-apoptotic factor C/EBP homologous protein (Chop, encoded by DNA damage-inducible transcript 3 [Ddit3]) (Costa-Mattioli and Walter 2020, Vasudevan, Neuman et al. 2020). Although Chop is traditionally associated with apoptosis, its transcriptional targets exhibit significant overlap with those of Atf4, encompassing genes that paradoxically support cell survival and growth (Marciniak, Yun et al. 2004, Han, Backa et al. 2013, Uzi, Barda et al. 2013). These findings underscore the context-dependent duality of the Atf4-Chop axis across different stress conditions, highlighting the need to elucidate its precise regulatory roles in specific physiological settings.

Through spatial transcriptomics (ST) and functional analyses, we demonstrate that zonal heterogeneity in APAP metabolism drives a stress-adaptive response in mid-zone hepatocytes during early APAP-induced liver injury. This response is mediated by the Atf4-Chop-Btg2 axis, which orchestrates a transient proliferation arrest to prioritize cellular survival over regenerative capacity. Our findings elucidate the spatial and molecular mechanisms governing hepatocyte adaptation to acute injury, uncovering a critical survival-regeneration trade-off in early phase of acute liver injury.

## Results

### Mid-zone hepatocytes exhibit the most pronounced proliferative decline during early acute liver injury

To assess hepatocyte proliferation during early APAP-induced liver injury, wild-type C57BL/6J mice received an intraperitoneal injection of 300 mg/kg APAP, and liver sections were analyzed at 0, 3, 6, 12, and 24 hours post-injection. Immunohistochemical staining for the proliferation marker Ki67 revealed that under homeostatic conditions (0 h), mid-zone hepatocytes exhibited approximately three-fold higher basal proliferative activity than periportal (PP) and pericentral (PC) hepatocytes, consistent with previous reports (Wei, Wang et al. 2021). During the early injury phase (3–12 h), the number of Ki67⁺mid-zone hepatocytes declined sharply from roughly 70 to 20 per section, reaching levels comparable to those observed in the PP and PC zones (Figure 1A, B). By 24 h post-APAP, mid-zone proliferation gradually recovered toward baseline levels (Figure 1A, B). Although all zones exhibited reduced proliferation during the injury initiation phase, the mid-zone showed the most substantial decline (Figure 1C), as further corroborated by value-change analysis relative to the 0 h time point (Figure 1D). Collectively, these data identify the mid-zone as the region most profoundly affected by transient proliferative suppression during early APAP-induced acute liver injury.

**Figure 1.**
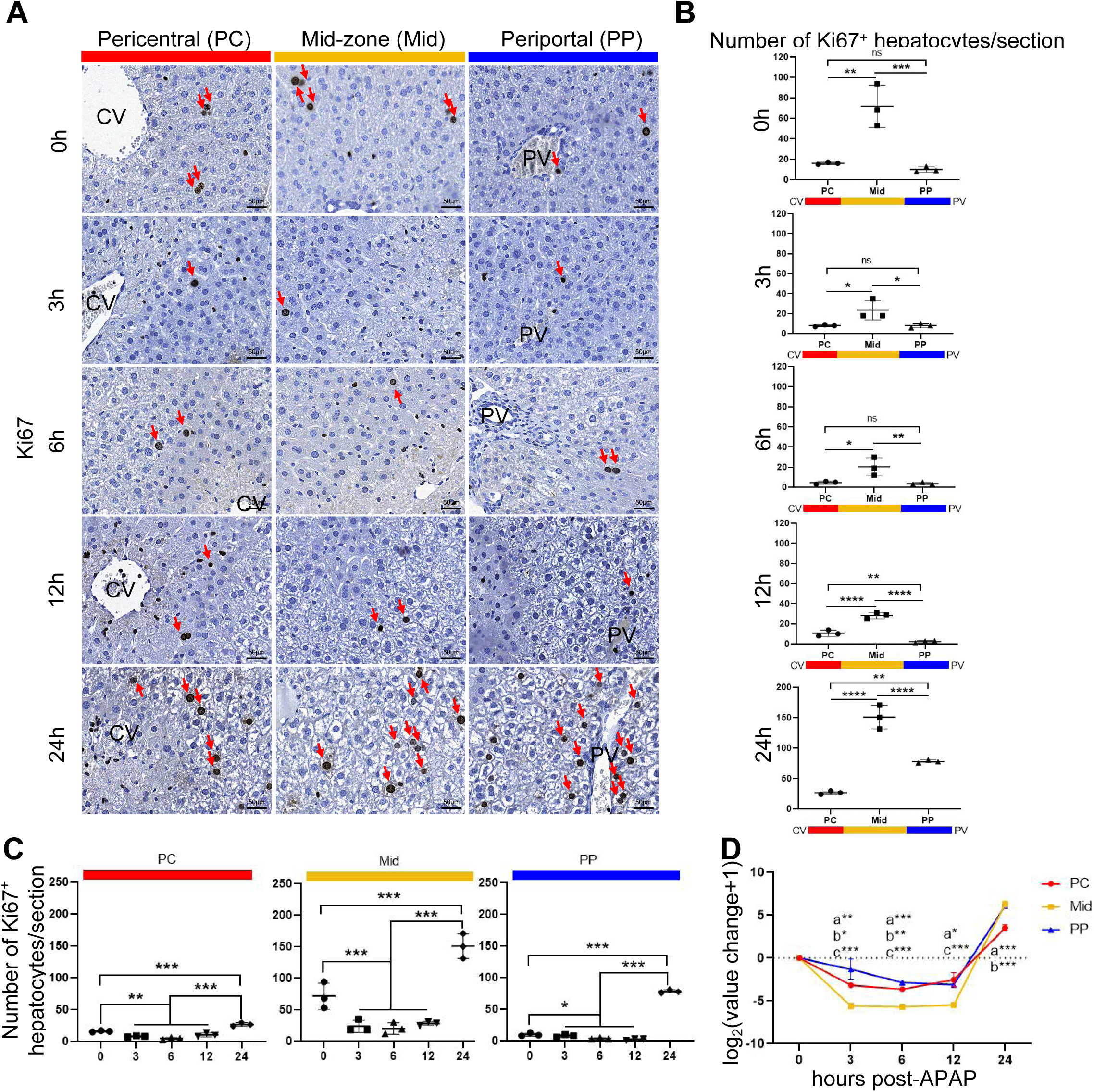
Mid-zone hepatocytes exhibit the most pronounced proliferative decline in early acute liver injury. Wildtype C57BL/6J mice were intraperitoneally injected with 300 mg/kg APAP for 0, 3, 6, 12 and 24 h. **(A)** Immunohistochemical staining of Ki67 to detect proliferating hepatocytes in liver sections at 0, 3, 6, 12 and 24 h post-APAP. Representative images of pericentral (PC), middle (Mid) and periportal (PP) are shown. Red arrows indicate Ki67+ hepatocytes. **(B)** Quantitative analysis of Ki67+ hepatocytes across liver zones (PC, Mid, PP) at 0, 3, 6, 12, and 24 h post-APAP. **(C)** Quantitative analysis of Ki67+ hepatocytes across 0, 3, 6, 12, and 24 h post-APAP in each liver zone (PC, Mid, PP). **(D)** Changes of Ki67+ hepatocytes across liver zones (PC, Mid, PP) at 0, 3, 6, 12 and 24 h post-APAP. Value change=number of Ki67+ hepatocytes at x h post-APAP - the number of Ki67+ hepatocytes at 0 h post-APAP. (a) denotes significance between PC and Mid regions, (b) denotes significance between PC and PP regions, and (c) denotes significance between Mid and PP regions. Data points represent mean hepatocyte counts from six high-power fields per zonal layer (n=3 mice/group) in B and C. Data are represented as means ± SD; One-way ANOVA (B,C,D).*p < 0.05; **p < 0.01; ***p < 0.001; ****p < 0.0001; ns, not significant.

### Mid-zone hepatocytes show distinct gene expression profiles

To investigate the mechanism underlying proliferation pause during early APAP-induced liver injury, liver sections from wild-type mice treated with APAP for various time points were subjected to ST using the Visium (10 × Genomics) platform (*Figure 2-figure supplement 1A*). Given inter-mouse variability, one representative sample per time point was selected based on H&E histology and serum ALT levels (*Figure 2-figure supplement 1B*). Under normal conditions (APAP 0h), unbiased transcriptional clustering partitioned each lobule into three distinct zones: PP (*Alb, Mup20, Cyp2f2*), PC (*Glul, Cyp2e1, Cyp1a2*), and mid-zone (intermediate between PP and PC, with elevated *Igfbp2* and *Hamp*) (Figure 2A, *Figure 2-figure supplement 1C*). Following APAP-induced injury (3h, 6h), the PC zone remained identifiable by residual PC signature genes despite necrosis and reduced transcription, while PP genes remained largely unchanged. The Mid zone was distinguishable as the intervening cluster exhibiting marked transcriptional reprogramming (*e.g.*, *Sqstm1, Igfbp1*) (Figure 2A and *Figure 2-figure supplement 1C*). UMAP projections and spatial heatmaps of *Glul* and *Cyp2f2* confirmed persistent, though dynamic, zonal clustering across all time points (*Figure 2-figure supplement 1D--F*). These data indicate that the fundamental lobular architecture remains spatially resolved during the early injury phase, enabling zone-specific transcriptional analysis.

**Figure 2.**
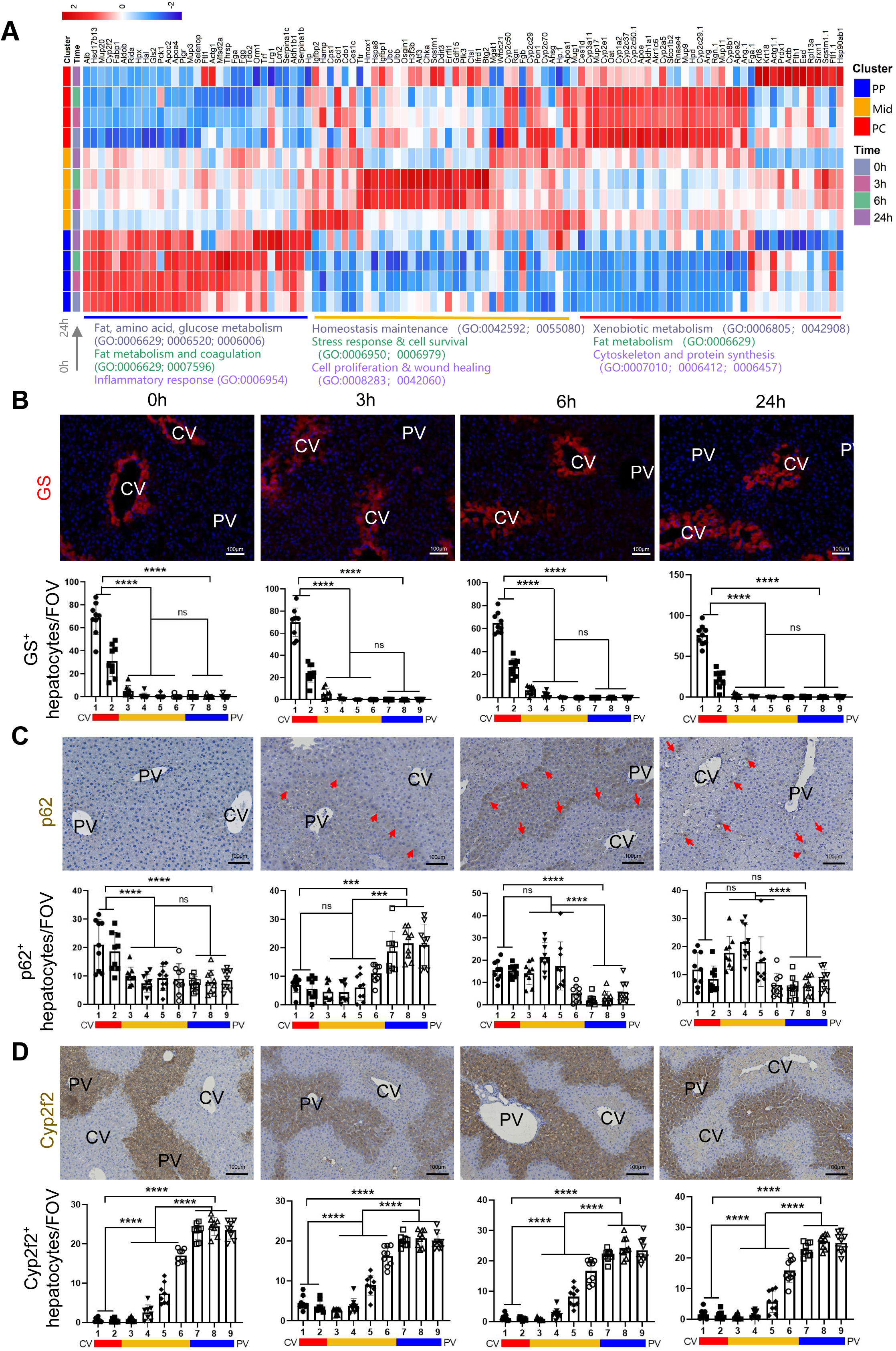
Mid-zone hepatocytes show distinct gene expression profiles. **(A)** Heatmap of the top 10 differentially expressed genes in periportal (PP), midzonal (Mid) and pericentral (PC) hepatocyte zones at 0, 3, 6 and 24 h post-APAP. To highlight relative expression shifts across zones, this heatmap is visualized using row-wise Z-score standardization, which scales each gene to a uniform range and emphasizes spatial switching patterns rather than absolute transcript abundance. Functional categories at the bottom were assigned using Gene Ontology (GO) terms (biological process, cellular component, molecular function); for genes annotated with multiple GO terms, the most representative or significantly enriched term was selected. **(B)** Immunofluorescence staining for glutamine synthetase (GS, a pericentral marker) with DAPI (blue) in liver sections at 0, 3, 6, and 24 h after APAP administration. Quantification of the zonal distribution of GS-positive cells at the indicated time points. **(C)** Immunohistochemical detection of p62 (Sqstm1 protein, a midzone marker during early injury) in liver sections at 0, 3, 6, and 24 h after APAP administration. Red arrows indicate representative p62-positive hepatocytes. Quantification of the zonal distribution of p62-positive cells at the indicated time points. **(D)** Immunohistochemical staining of Cyp2f2 (a periportal marker) in liver sections at 0, 3, 6, and 24 h after APAP administration. Quantification of the zonal distribution of Cyp2f2-positive cells at the indicated time points. Data are presented as mean ± SD. One-way ANOVA (B-D). *p < 0.05, **p < 0.01, ***p < 0.001, ****p < 0.0001; ns, not significant.

Differential expression analysis across zones and time points revealed distinct zonal trajectories (Figure 2A). In the PP zone, gene expression remained relatively stable across 0–6 h, with inflammatory genes (*Orm1, Serpina1c*) rising only at 24 h. The mid-zone exhibited a clear temporal progression. At 0 h, it expressed proliferative (*Igfbp2*), metabolic (*Scd1, Cdo1*), and homeostatic (*Hamp, Cps1*) genes. By 3–6 h, these were replaced by stress-response and autophagy genes (*Hmox1, Hspa8, Atf3, Ubc, Sqstm1*). By 24 h, wound-healing genes (*Fgg, Fgl1*) became prominent. The PC zone showed early downregulation of xenobiotic metabolism (*Cyp2e1, Cyp1a2*), followed by upregulation of cytoskeletal (*Actb, Krt8*) and redox (*Prdx1, Srxn1*) genes at 24 h. This temporal trajectory identifies the mid-zone as the most transcriptionally dynamic region, undergoing a functional switch from homeostasis to active stress adaptation during the initial injury phase.

Focusing on the mid-zone at 6 h versus 0 h, volcano plot analysis identified significant upregulation of stress-related genes (*Atf3, Hmox1, Ddit3, Atf4*) alongside cell-cycle and apoptosis regulators (*Btg2, Egr1, Sqstm1*) (*Figure 2-figure supplement 1G*). Among these, we selected *Sqstm1* (encoding p62) for validation as a representative stress-response gene with previously characterized roles in autophagy and oxidative stress. To validate our zonation assignment, we compared ST clusters with a nine-layered radial classification from the central vein (CV) to the portal vein (PV). Immunostaining for GS (PC), p62 (mid-zone), and Cyp2f2 (PP) confirmed that our PC, Mid, and PP zones corresponded to layers 1–2, 3–6, and 7–9, respectively (Figure 2B--D). These spatial correlations confirm that the mid-zone is the primary site of early stress-response gene induction, with p62 serving as a robust regional marker.

ST-based analysis of Ki67+ spots recapitulated the transient proliferative reduction at 3-6 h (*Figure 2-figure supplement 1I*). To further dissect cell-cycle dynamics, we assessed S-phase and G2/M-phase gene signatures. Mid-zone hepatocytes displayed a transient increase in S-phase scores at 3 h, followed by a decline at 6 and 24 h; conversely, G2/M scores were reduced at 3-6 h and returned to baseline by 24 h (*Figure 2-figure supplement 1J*). Collectively, these ST analyses establish that mid-zone hepatocytes adopt a spatially confined stress-adaptive transcriptional program—marked by p62 upregulation and a impaired cell-cycle progression—during the early phase of APAP-induced injury, providing a molecular framework for the proliferative pause observed histologically.

### Zonal metabolism of APAP leads to stress response in Mid-zone hepatocytes

To determine what triggers the mid-zone stress response, we first examined the spatial distribution of APAP-metabolizing cytochrome P450 (Cyp) enzymes. Cyp enzymes metabolize APAP to NAPQI, which causes oxidative stress and cellular damage unless neutralized by GSH (Figure 3A). ST analysis revealed a zonal gradient in the expression of most Cyp transcripts, with highest levels in the PC zone progressively decreasing toward the PP zone (Figure 3B and *Figure 3-figure supplement 1A*). Immunohistochemical staining for Cyp1a2 confirmed this gradient at the protein level under homeostatic conditions (0 h), with strong pericentral signal diminishing toward the portal tracts. However, following APAP treatment (3–24 h), Cyp1a2 expression was largely lost within the necrotic CV region, while the adjacent mid-zone remained the primary area retaining detectable Cyp1a2 signal (Figure 3C and 3D). This spatial shift suggests that selective necrosis of Cyp-rich pericentral hepatocytes effectively relocates the residual metabolic capacity to the surviving mid-zone parenchyma (Figure 3E).

**Figure 3.**
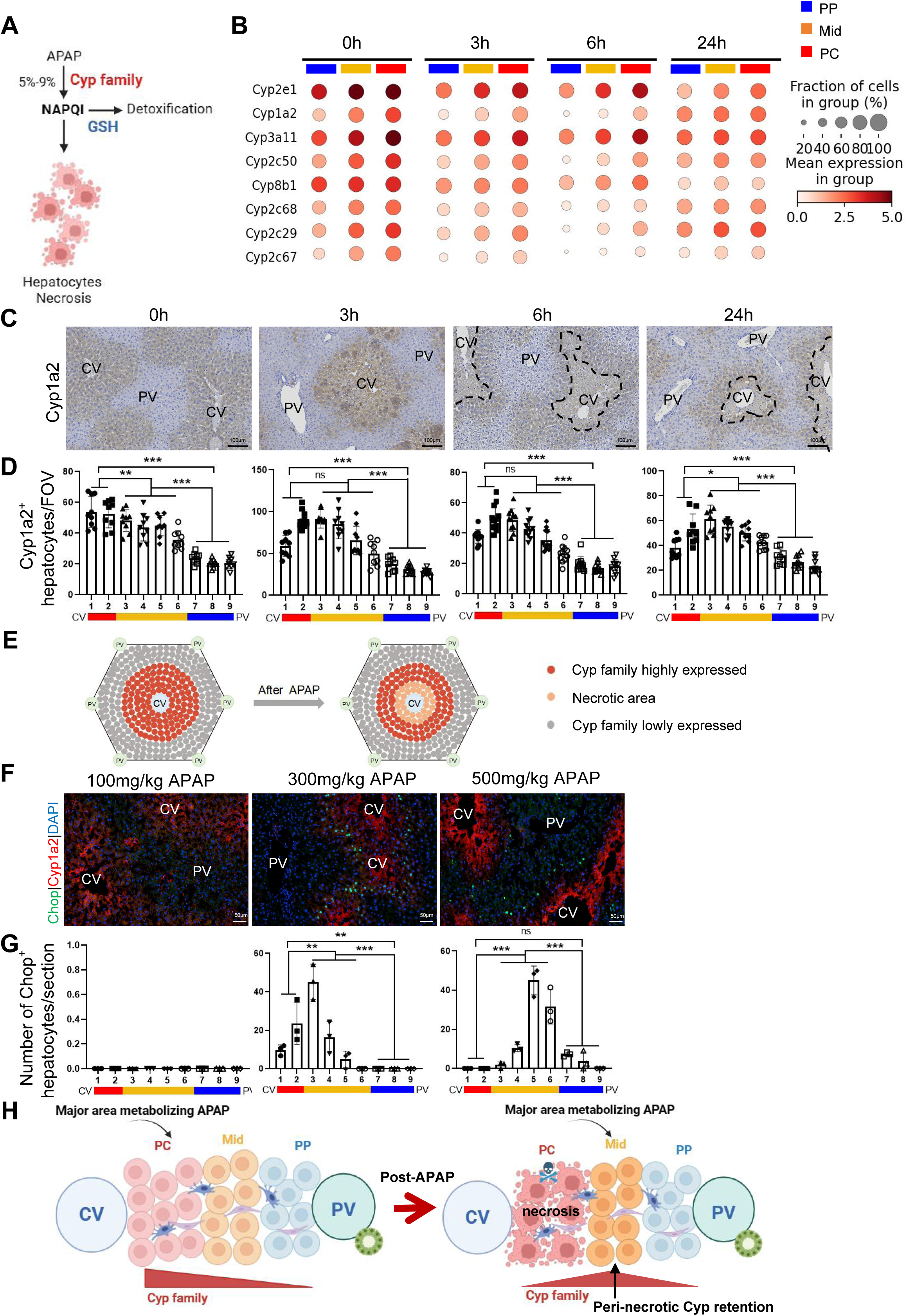
Zonal metabolism of APAP leads to stress response in Mid-zone hepatocytes. **(A)** Schematic figure illustrating the metabolic process of liver injury caused by APAP. **(B)** Dot plot displays the log-normalized absolute transcript abundance of the Cyp family in hepatocyte zones (zones PP, Mid, and PC) at 0, 3, 6, 24 h post-APAP. **(C)** Immunohistochemical staining of Cyp1a2 to detect its dynamic changes in hepatocyte zones (from CV to PV) at 0, 3, 6, 24 h post-APAP. Injured area is outline by black dashed lines. **(D)** Quantification of zonal distribution of Cyp1a2-positive cells in liver sections at 0, 3, 6, and 24 h post-APAP, respectively. **(E)** Schematic figure illustrating expression changes of Cyp1a2 before and after APAP. **(F)** Immunofluorescence staining of Chop protein (green) to observe the dynamic changes in its expression across hepatocyte zones (from CV to PV) at 6 h post-APAP with doses of 100, 300, or 500 mg/kg. Cyp1a2 protein (red) staining highlights the area around the CV. Cell nuclei are stained with DAPI (blue). **(G)** Quantification of zonal distribution of Chop-positive cells in liver sections from mice treated with varying doses of APAP– 100, 300, or 500 mg/kg. **(H)** Schematic explaining why mid-lobular hepatocytes display a distinct gene expression program early after APAP injury. Cyp enzymes are expressed in a decreasing gradient from the central vein (CV) to the portal vein (PV). APAP is therefore metabolized primarily by pericentral Cyp-expressing hepatocytes; their necrosis shifts the main site of residual Cyp activity to the mid-lobular zone. High local APAP metabolism then drives a specific stress-response program, accounting for the unique early transcriptional signature of the mid-zone. Data are represented as means ± SD; One-way ANOVA (D,G). *p < 0.05; **p < 0.01; ***p < 0.001; ****p < 0.0001; ns, not significant.

To test whether the extent of necrosis dictates the magnitude and localization of the mid-zone stress response, we administered escalating APAP doses (100, 300, and 500 mg/kg). H&E staining confirmed a dose-dependent expansion of necrotic areas (*Figure 3-figure supplement 1B--C*). We then assessed Chop (the protein product of Ddit3), a key mid-zone stress marker whose transcript was upregulated at 6 h post-APAP (Figure 2B). Chop immunoreactivity was absent at the lowest dose, where necrosis was minimal. As the dose increased and the necrotic area expanded, Chop expression emerged immediately adjacent to the necrotic front and progressively shifted toward the portal veins, always surrounding the injured territory (Figure 3F and 3G). Thus, CV necrosis establishes the adjacent peri-necrotic region as the functional epicenter of residual APAP metabolism, directly linking zonal Cyp distribution to localized stress response (Figure 3H).

### The Atf4-Chop axis emerges as pivotal in Mid-zone during early acute injury

To identify upstream drivers of the mid-zonal stress program, we constructed gene regulatory networks (GRNs) from the ST data and ranked the top 10 regulons per zone at 3 and 6 hours post-APAP (Figure 4A and *Figure 4-figure supplement 1A*). GRN analysis revealed simultaneous activation of several transcription factors (TFs) in the mid-zone across both time points, including Atf4, Fos, Nfil3, Egr1, Maff, Atf3, and Ddit3. These TFs include known stress-response regulators (*Atf4, Atf3, Ddit3*), immediate-early genes (*Fos, Egr1*), and oxidative/metabolic regulators (*Maff, Nfil3*), suggesting a multi-faceted adaptive program (Ohoka, Yoshii et al. 2005, Woo, Cui et al. 2009, von Scheidt, Zhao et al. 2021, Bunch, Kim et al. 2023, Lin, Hsu et al. 2024). Among the top 10 mid-zone regulons, Ddit3 exhibited the highest inferred activity, while Atf4 ranked seventh (Figure 4B and *Figure 4-figure supplement 1B*). Since the Atf4-Ddit3 axis is a well-established regulator of stress adaptation (Zhang, Guo et al. 2022), the coordinated activation of this transcriptional network prompted us to investigate this pathway further. Violin plots confirmed prominent mid-zone enrichment of both Atf4 and Ddit3 at 3 and 6 hours (Figure 4C and *Figure 4-figure supplement 1C*). GRNs illustrated connectivity between Atf4 and its targets (*e.g.*, *Gdf15, Hspa8, Ddit3*) as well as between Ddit3 and its targets (*e.g.*, *Hspa8, Hmox1, Osgin1, Gadd45a, Egr1*) (*Figure 4-figure supplement 1D*). Heatmaps further demonstrated that the target genes of both TFs were selectively expressed in mid-zone hepatocytes at 6 hours post-APAP (Figure 4D), and many of these targets overlapped with the mid-zone DEGs identified earlier. This coordinated enrichment and target overlap strongly implicate the Atf4-Ddit3 axis as a principal driver of the mid-zone transcriptional reprogramming.

**Figure 4.**
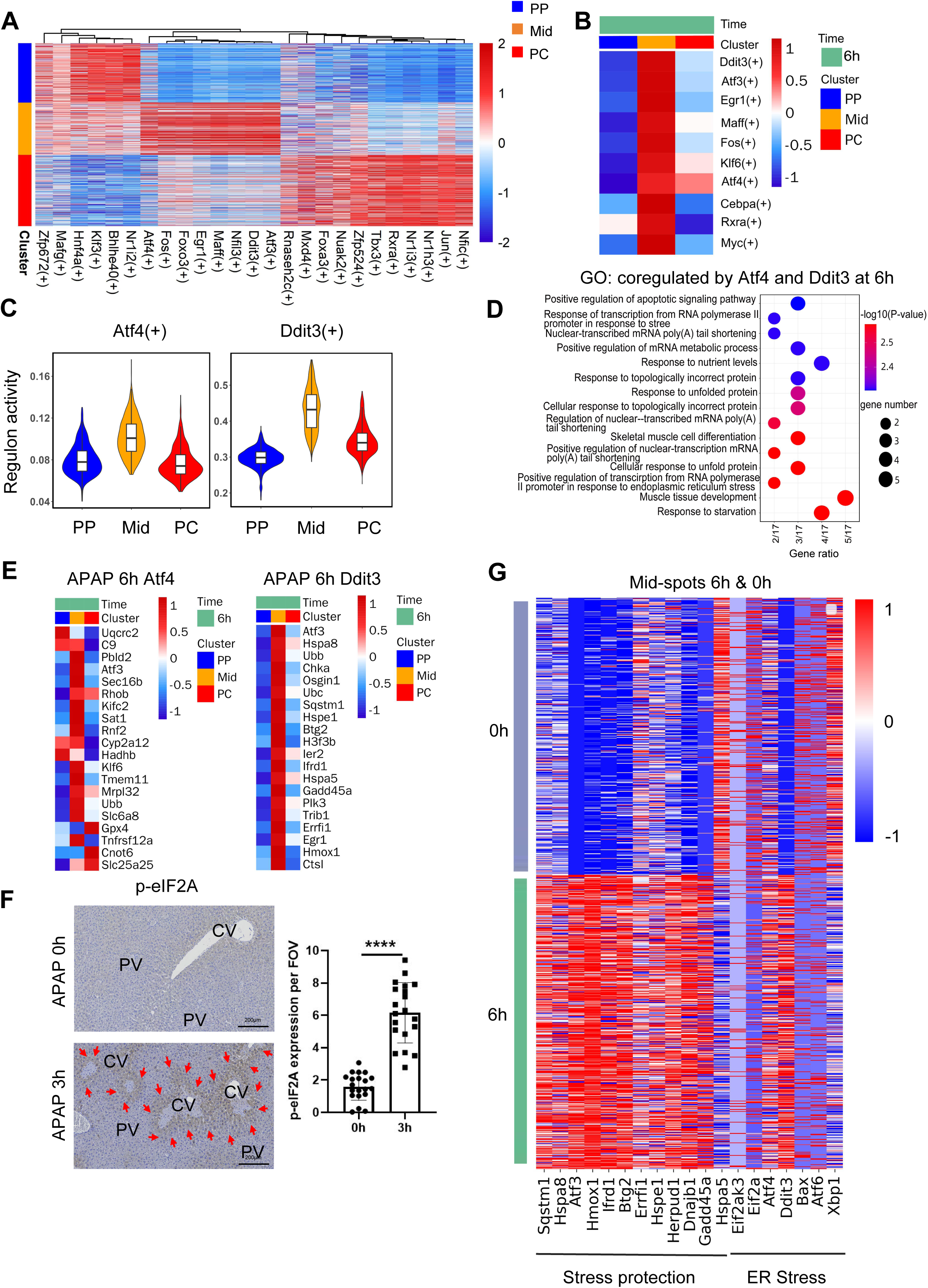
The Atf4-Ddit3 axis emerges as pivotal in Mid-zone during early acute injury. **(A)** Heatmap displays the area under the curve (AUC) scores of transcription factor (TF) motifs, estimated per dot by Single-Cell Regulatory Network Inference and Clustering (SCENIC), highlighting gene regulatory networks and differentially activated TF motifs in hepatocyte zones (PP, Mid, PC) at 6 h post-APAP. Columns represent TF motifs, rows represent dots, and color intensity indicates AUC scores. **(B)** Heatmap shows inferred transcription factors (TFs) activity across different zonal regions at 6 h post-APAP. Activity was quantified as regulon enrichment scores using AUCell. The color scale represents regulon activity scores. **(C)** Violin plots shows regulon activity of Atf4 and Ddit3 in each zonation at 6 h post-APAP. Activity was quantified as regulon enrichment scores using AUCell. **(D)** GO pathway analysis reveals the top 15 enriched pathways for genes co-regulated by Atf4 and Ddit3 in mid hepatocyte zones at 6 h post-APAP. **(E)** Heatmap showing the target genes expression of Atf4 (left) and Ddit3 (right) in hepatocyte zones (zones PP, Mid, and PC) at 6h post-APAP. **(F)** Immunohistochemistry staining of p-eIF2A in liver sections from mice at 0 h and 3 h post-APAP. p-eIF2A-positive hepatocytes are indicated by red arrows. The number of p-eIF2A-positive cells per field of view (FOV) is quantified. n=3 mice/group. **(G)** Heatmap displays stress response and ER stress-related genes from differentially expressed genes (DEGs) identified in Mid hepatocyte zones at 0 and 6 h post-APAP. Data are represented as means ± SD; Unpaired Student’s t-test (F). *p < 0.05; **p < 0.01; ***p < 0.001; ****p < 0.0001; ns, not significant.

GO analysis of Atf4–Ddit3 co-regulated targets revealed enrichment for pathways related to unfolded protein response, cellular stress, autophagy, and apoptotic signaling (Figure 4E and *Figure 4-figure supplement 1E--F*). Immunohistochemical staining confirmed increased phosphorylation of eIF2α (p-eIF2α) in the mid-zone at 3 hours post-APAP, indicating ISR activation (Figure 4F).However, when we examined canonical ER-stress markers, we found that genes such as Eif2ak3, Atf6, and Xbp1 either remained unchanged or were downregulated at 3 and 6 hours post-APAP, despite strong Atf4 and Ddit3 upregulation (Figure 4G and *Figure 4-figure supplement 1G*). These data indicate that the mid-zone stress response engages the ISR through eIF2α phosphorylation but does not constitute a classical, full-scale ER stress/unfolded protein response. Instead, the response appears selectively routed through the Atf4–Ddit3 branch.

Reanalysis of a public APAP-injury scRNA-seq dataset (Matchett, Wilson-Kanamori et al. 2024). confirmed zonal hepatocyte clustering (PP, Mid, PC) at 3 and 6 hours post-APAP, with mid-zone enrichment of Hmox1, Atf3, and Ubc (*Figure 4-figure supplement 2A--C*). GO analysis of mid-zone upregulated genes highlighted stress-response and cell-death processes (*Figure 4-figure supplement 2D--E*). SCENIC analysis independently identified activation of Ddit3 and Atf3 motifs in the mid-zone (*Figure 4-figure supplement 2F--G*).This external dataset validates our ST-based observations and reinforces the specificity of the mid-zone stress-regulatory program.

Analysis of human snRNA-seq data from healthy and APAP-induced acute liver failure (ALF) explants (GSE223561) revealed distinct periportal (*ALB, CYP2F2*), mid-zonal (*IGFBP2, HAMP*), and pericentral (*GLUL, CYP2E1*) hepatocyte populations (*Figure 4-figure supplement 3A-D*). In healthy human livers, mid-zonal hepatocytes showed detectable expression of stress regulators (*DDIT3, ATF4, HMOX1*) alongside low cell-cycle gene expression. In APAP patients, mid-zonal stress regulators were further elevated; however, in contrast to our early murine model, proliferation-associated genes (*MKI67, CCNB1, CDK1*) were also upregulated in the same mid-zonal population (*Figure 4-figure supplement 3E*). Human ST data from two APAP patients revealed inter-patient heterogeneity: one specimen showed stress-gene enrichment with reduced proliferation markers in the peri-necrotic mid-zone, while the other displayed a predominantly proliferative transcriptional profile with weaker stress signaling (*Figure 4-figure supplement 3F*). These human data suggest that while the mid-zonal stress program is conserved, the balance between stress adaptation and proliferation is dynamic and likely stage-dependent, consistent with the temporal progression observed in our mouse model.

### The Atf4-Ddit3 axis protects hepatocytes from liver injury

To assess the functional relevance of the Atf4–Ddit3 axis, we first examined endogenous Atf4 protein dynamics. Immunohistochemical staining revealed progressive nuclear Atf4 accumulation in mid-zone hepatocytes between 0 and 6 hours post-APAP (Figure 5A). This temporal and spatial pattern aligns with the transcriptional upregulation observed in our ST data, confirming mid-zone-specific Atf4 activation.

**Figure 5.**
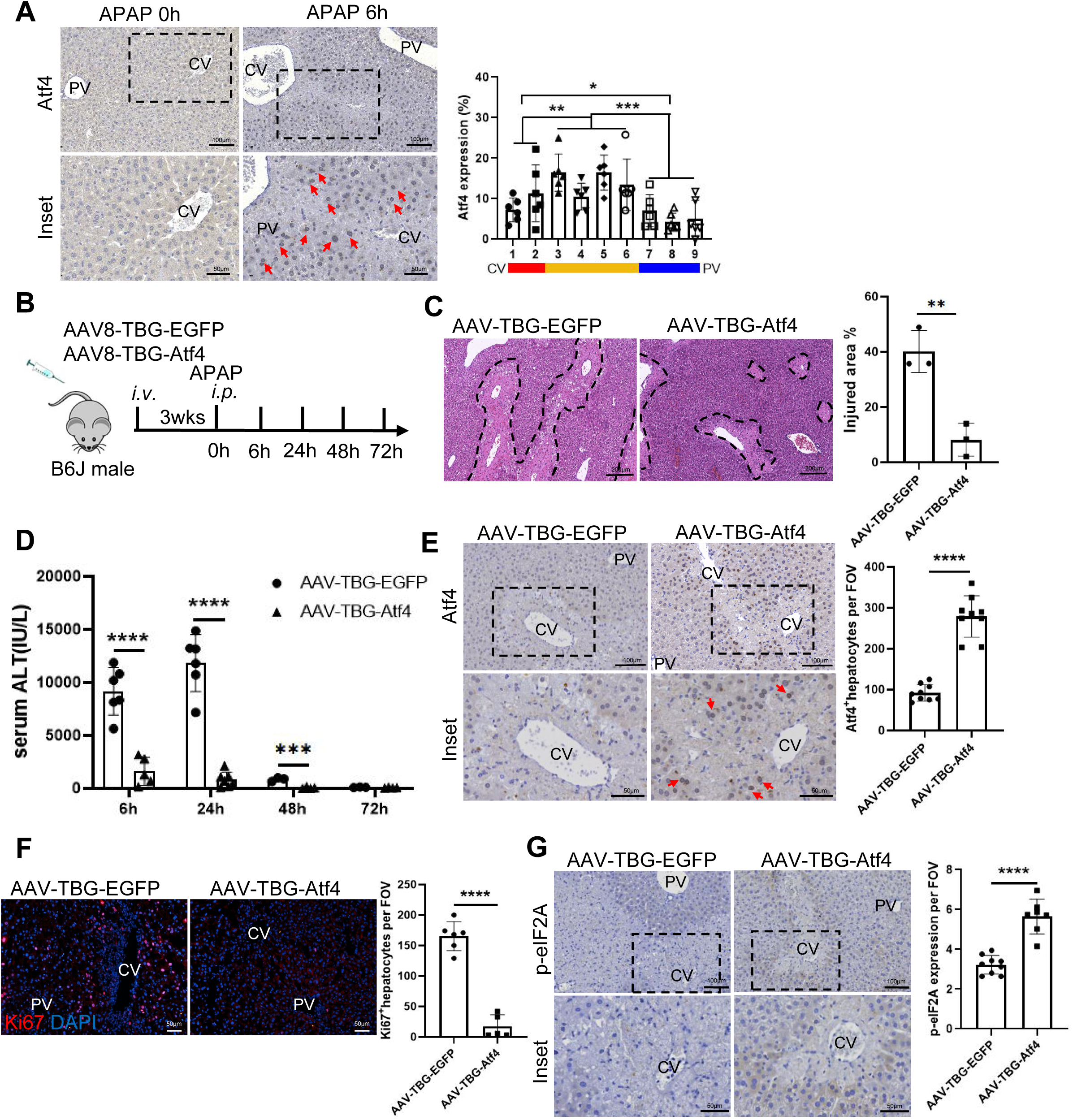
The Atf4-Ddit3 axis protects hepatocytes from liver injury. **(A)** Immunohistochemical staining of Atf4 in liver sections at 0 and 6 h post-APAP. Red arrows indicate Atf4-positive hepatocytes. Zonal distribution of Atf4-positive cells in liver sections at 6 h post-APAP is quantified. The statistic is the percentage of Atf4-positive hepatocytes in each layer over the total number of Atf4-positive hepatocytes. n=3 mice. **(B)** Schematic figure illustrating the overexpression of Atf4 via AAV in hepatocytes of wildtype C57BL/6J mice. Overexpression of EGFP is used as a control. **(C)** H&E staining showing liver morphology from mice overexpressing AAV-TBG-EGFP or AAV-TBG-Atf4 at 6 h post-APAP. The injured area is outlined by black dashed lines. The percentage of injured area is quantified. n=3 mice/group. Necrotic areas outlined by loss of cellular architecture on H&E. **(D)** Serum levels of ALT is measured in mice overexpressing AAV-TBG-EGFP or AAV-TBG-Atf4 at 6, 24, 48, and 72 h post-APAP. n=6 mice/group. **(E)** Immunohistochemistry staining of Atf4 in liver sections from mice overexpressing AAV-TBG-EGFP or AAV-TBG-Atf4 at 24 h post-APAP. Atf4-positive hepatocytes are indicated by red arrows. The number of Atf4-positive cells per field of view (FOV) is quantified. n=9 mice/group. (F) Immunofluorescence staining of Ki67 (red) was performed to assess proliferating hepatocytes in mice overexpressing AAV-TBG-EGFP or AAV-TBG-Atf4 at 72 h post-APAP. Cell nuclei were stained with DAPI (blue). The number of Ki67-positive cells per FOV is quantified. n=6 mice/group. **(G)** Immunohistochemistry staining of p-eIF2A in liver sections from mice overexpressing AAV-TBG-EGFP or AAV-TBG-Atf4 at 24 h post-APAP. p-eIF2A-positive hepatocytes are indicated by red arrows. The number of p-eIF2A-positive cells per FOV is quantified. n=6 mice/group. Data are represented as means ± SD; Unpaired Student’s t-test (C-G). One-way ANOVA (A). *p < 0.05; **p < 0.01; ***p < 0.001; ****p < 0.0001; ns, not significant.

To test causality, we overexpressed Atf4 specifically in hepatocytes using adeno-associated virus 8 (AAV8) - thyroxine-binding globulin (TBG) (Figure 5B). Atf4 overexpression markedly attenuated APAP-induced liver injury, as evidenced by reduced necrotic areas on H&E staining and significantly lower serum ALT levels (Figure 5C and 5D). Immunohistochemical staining confirmed nuclear Atf4 localization in hepatocytes adjacent to injured regions (Figure 5E). Furthermore, Atf4 overexpression resulted in decreased Ki67⁺proliferating cells and increased p-eIF2α immunoreactivity (Figure 5F and 5G). Together, these data demonstrate that elevated Atf4 actively promotes hepatocyte survival by enhancing ISR signaling, suppressing proliferation, and mitigating early liver damage.

### The Atf4-Ddit3 axis protects the liver by pausing proliferation via Btg2

Atf4 and Ddit3 are key transcriptional regulators of ISR, and their downstream target genes converge on overlapping stress-responsive pathways (Ohoka, Yoshii et al. 2005, Woo, Cui et al. 2009, Kaspar, Oertlin et al. 2021). To dissect the downstream effectors of the Atf4–Ddit3 axis, we mapped Ddit3 genomic occupancy by Cut&Run at 6 h post-APAP—the peak of mid-zone Ddit3 activity during the injury-initiation phase. Compared with input control, we detected robust Ddit3-binding signals across chromatin domains (Figure 6A). Overlapping 6,157 Ddit3-bound genes with the 47 mid-zone–specific DEGs identified 39 overlapping targets, termed Ddit3-binding DEGs (Figure 6B). IGV tracks confirmed Ddit3 occupancy at regulatory elements of Btg2, Atf3, Dnajb1, and Egr1 (Figure 6C).

**Figure 6.**
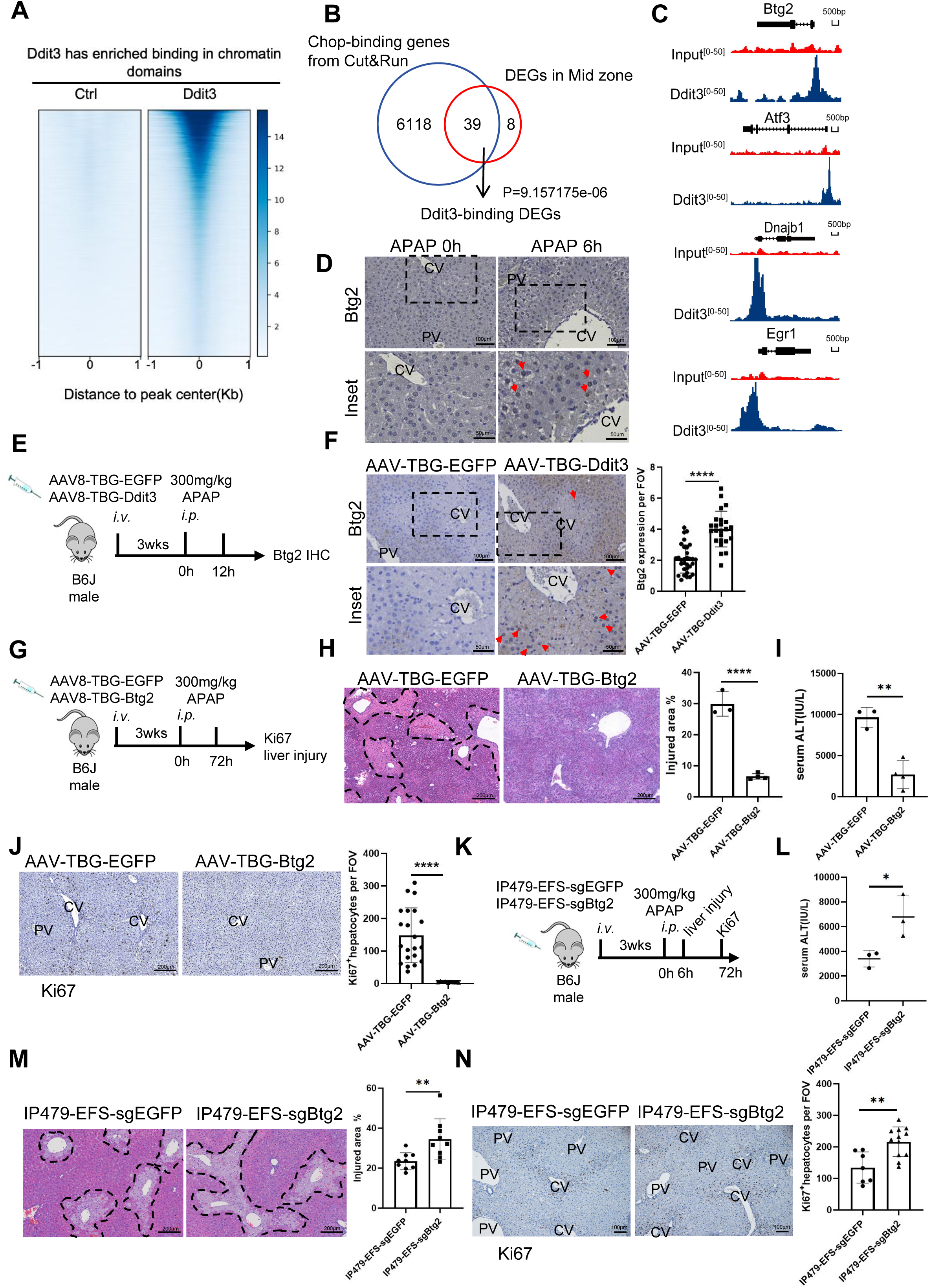
The Atf4-Ddit3 axis protects the liver by pausing proliferation via Btg2. **(A)** Heatmap of Ddit3-binding signals in chromatin domains. **(B)** Overlap of Ddit3-binding genes by Cut&Run with differentially expressed genes (DEGs) in mid zone was shown. These genes were referred as Ddit3-binding DEGs. **(C)** Integrative Genomics Viewer (IGV) plot shows the Cut&Run peaks of Ddit3 and input on Btg2-site, Ifrd1-site, Fos-site and Egr1-site. **(D)** Immunohistochemical staining of Btg2 in liver sections at 0 and 6 h post-APAP. Red arrows indicate Btg2-positive hepatocytes. **(E)** Schematic figure illustrating overexpression of Ddit3 via AAV in hepatocytes of wildtype C57BL/6J mice. Overexpression of EGFP is used as a control. **(F)** Immunohistochemical staining of Btg2 in liver sections from mice overexpressing either AAV-TBG-EGFP or AAV-TBG-Ddit3 at 12 h post-APAP. Btg2-positive hepatocytes are indicated by red arrows. Btg2-positive hepatocytes in liver sections is quantified. n=4 mice/group. **(G)** Schematic figure illustrating overexpression of Btg2 via AAV in hepatocytes of wildtype C57BL/6J mice. Overexpression of EGFP is used as a control. **(H)** H&E staining shows morphology of livers from AAV-TBG-EGFP or AAV-TBG-Btg2 overexpressing mice at 6 h post-APAP. Injured area is outlined by black dashed lines. The percentage of injury area is quantified. n=3 mice/group. Necrotic areas outlined by loss of cellular architecture on H&E. **(I)** Serum levels of ALT is measured at 6h post-APAP in AAV-TBG-EGFP or AAV-TBG-Btg2 overexpressing mice. n=3 mice/group. **(J)** Immunohistochemical staining of Ki67 in liver sections from mice overexpressing either AAV-TBG-EGFP or AAV-TBG-Btg2 at 72 h post-APAP. Ki67-positive hepatocytes are indicated by red arrows. Ki67-positive cells in liver sections is quantified. n=4 mice/group. **(K)** Schematic figure illustrating knockdown of Btg2 via AAV in hepatocytes of wildtype C57BL/6J mice. IP479-EFS-sgEGFP is used as a control. **(L)** Serum levels of ALT is measured at 6h post-APAP in IP479-EFS-sgEGFP or IP479-EFS-sgBtg2 knockdown mice. n=3 mice/group. **(M)** H&E staining shows morphology of livers from IP479-EFS-sgEGFP or IP479-EFS-sgBtg2 knockdown mice at 6 h post-APAP. Injured area is outlined by black dashed lines. The percentage of injury area is quantified. n=3 mice/group. Necrotic areas outlined by loss of cellular architecture on H&E. **(N)** Immunohistochemical staining of Ki67 in liver sections from IP479-EFS-sgEGFP or IP479-EFS-sgBtg2 knockdown mice at 72 h post-APAP. Ki67-positive cells in liver sections is quantified. n=3 mice/group. Data are represented as means ± SD; Unpaired Student’s t-test (F, H-N). **p < 0.01; ***p < 0.001; ****p < 0.0001; ns, not significant.

Among the Ddit3-binding DEGs, B-cell translocation gene 2 (Btg2) is cell cycle inhibitor (Park, Kim et al. 2008, Stupfler, Birck et al. 2016, Yuniati, Scheijen et al. 2019). Moreover, Btg2 is also coregulated by Ddit3 and Atf4. Immunohistochemical staining further revealed increased Btg2 protein expression around the injured area at 6 h post-APAP (Figure 6D). To determine whether Ddit3 drives Btg2 expression, we overexpressed Ddit3 in hepatocytes via AAV8-TBG (Figure 6E). Ddit3 overexpression significantly elevated Btg2 protein levels at 12 h post-APAP compared to EGFP controls (Figure 6F). We next overexpressed Btg2 directly (Figure 6G). Btg2 overexpression significantly reduced APAP-induced injury, as shown by decreased necrotic areas and serum ALT levels (Figure 6H, 6I), and markedly reduced Ki67⁺ hepatocyte numbers (Figure 6J).

To rule out potential confounds associated with AAV transduction, we measured Cyp2e1 protein levels, which showed only a modest decrease in both Atf4- and Btg2-overexpressing groups—well below the threshold known to confer significant protection (*Figure 6-figure supplement 1A*) (Ganetsky, Berg et al. 2019, Ye, Chen et al. 2022). AAV transduction efficiencies averaged 32% (EGFP), 18% (Atf4), and 23% (Btg2), with individual efficiency negatively correlating with ALT levels (*e.g.*, Pearson r = –0.7681, p = 0.0260 for Atf4; *Figure 6-figure supplement 1B--C*). These controls confirm that the protective effects are attributable to transgene expression rather than nonspecific Cyp2e1 inhibition. Forced Btg2 expression phenocopies the protective, proliferation-suppressive state observed during early injury, positioning Btg2 as a critical downstream mediator of the stress response.

To determine whether Btg2 is necessary for the protective arrest, we knocked down Btg2 using the AAV8-CasRx-sgRNA system (Figure 6K) (Zhou, Su et al. 2020). Despite a moderate (∼30%) reduction in Btg2 expression (*Figure 6-figure supplement 1D*), knockdown resulted in a significant ∼2-fold increase in serum ALT, ∼1.5-fold expansion of necrotic areas, and ∼1.8-fold increase in Ki67⁺hepatocytes compared to sgEGFP controls (Figure 6L--N). These data suggest that loss of Btg2 abrogates the protective brake on proliferation, sensitizing the liver to injury and permitting unscheduled cell-cycle activity under stress conditions.

To assess the broader applicability of this mechanism, we tested a carbon tetrachloride (CCl4)-induced acute liver injury model, which causes centrilobular necrosis through free-radical pathways rather than cytochrome P450–dependent metabolite formation (Wang, Wang et al. 2024). At 18 h post-CCl₄ injection, we observed elevated p-eIF2α, Atf4, Chop, and Btg2 levels alongside reduced Ki67 expression near injury sites (*Figure 6-figure supplement 2A--G*). This conservation across mechanistically distinct toxicants suggests that ISR-mediated Btg2 induction and subsequent proliferation arrest represent a general adaptive response to centrilobular injury. Collectively, these data establish a conserved functional cascade in which the Atf4-Ddit3 axis induces Btg2 to enforce transient proliferative restraint, prioritizing survival during early acute liver injury and preserving the capacity for subsequent regeneration.

## Discussion

Our study supports a model that during the early phase of APAP-induced liver injury, mid-zone hepatocytes undergo a transient proliferative restraint orchestrated by a distinctive transcriptional program centered on the Atf4–Chop–Btg2 axis. Spatial transcriptomics and functional assays demonstrate that this adaptive response promotes survival by upregulating the cell-cycle inhibitor Btg2, highlighting a trade-off between cytoprotection and proliferative capacity. Notably, zonal metabolic heterogeneity in APAP processing drives preferential activation of these stress pathways in mid-zone hepatocytes. These findings uncover a critical mechanism by which hepatocytes prioritize stress adaptation before initiating regeneration, offering new insights into the spatiotemporal regulation of liver repair (Figure 7).

**Figure 7.**
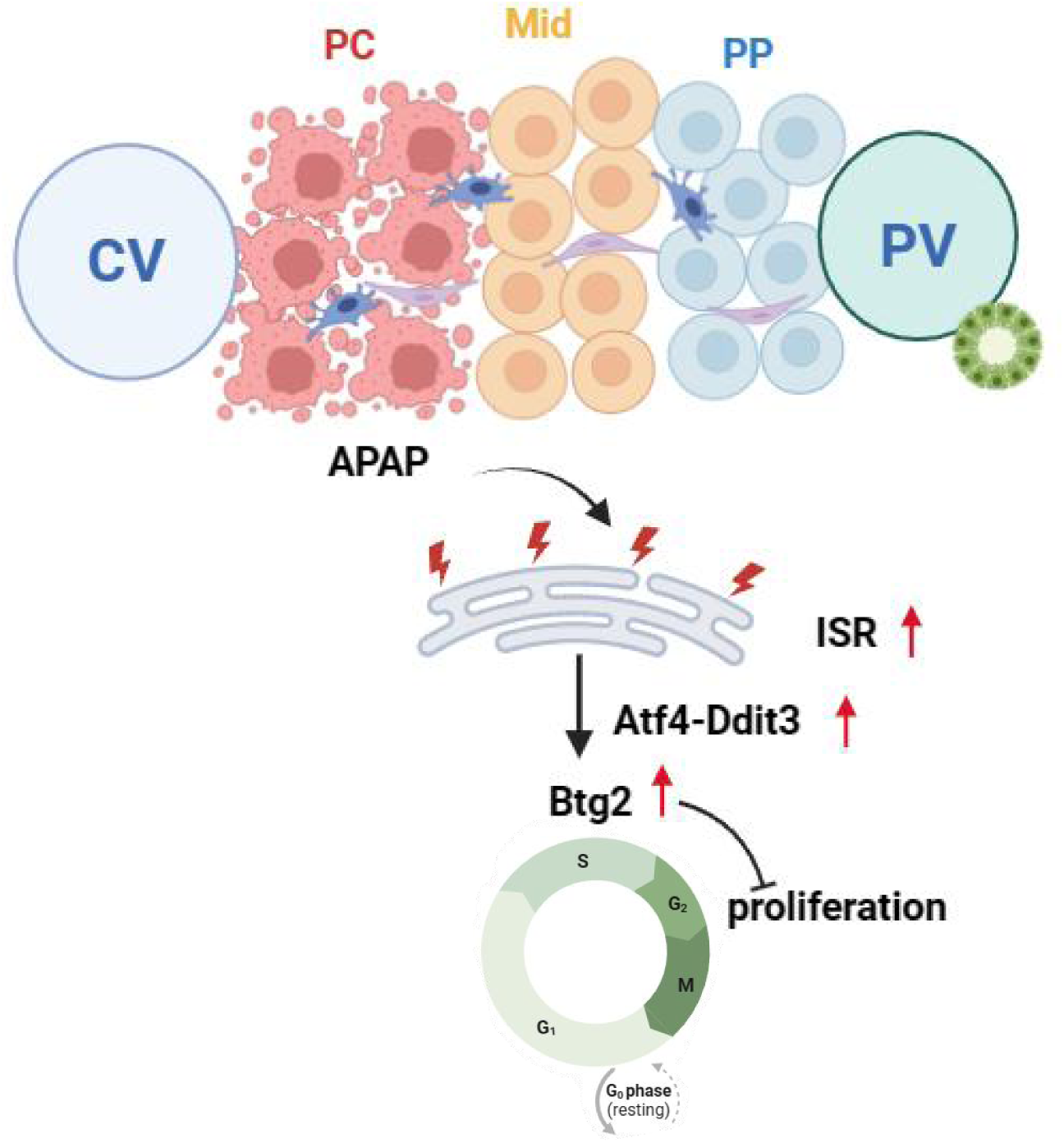
The Atf4-Ddit3 axis mediates integrated stress protection in mid-zone hepatocytes at the expense of proliferative capacity during early AILI. Schematic model depicting the adaptive response of mid-zone hepatocytes to acute liver injury, wherein activation of the Atf4-Ddit3 pathway upregulates Btg2 to induce transient cell cycle arrest, balancing survival and regeneration.

While the ISR shares upstream initiators with ER stress (*e.g.*, eIF2α phosphorylation), its outcomes are context-dependent. ER stress specifically activates the UPR through Ire1α, Atf6, and Perk (Wang, Thomas et al. 2006), with genetic studies demonstrating that knockout of Ire1α, Xbp1, or Ddit3 consistently attenuates liver damage, confirming the UPR’s contribution to hepatotoxicity (Nagy, Kardon et al. 2007, Hur, So et al. 2012, Uzi, Barda et al. 2013, Ye, Chen et al. 2022). Unlike classical ER stress, we find that mid-zone hepatocytes leverage the Atf4–Chop axis—defined here as a functional pathway of co-regulated transcriptional targets—to transiently halt proliferation via Btg2, favoring survival. This aligns with emerging evidence that Ddit3 can paradoxically support adaptation in mitochondrial stress (Kaspar, Oertlin et al. 2021, Kress, Jessen et al. 2023), suggesting that Ddit3 plays a context-dependent role in fine-tuning the ISR in mammals.

Our work resolves an early adaptive phase (3–12 h) of proliferation arrest that is temporally and functionally distinct from previously described zonal regeneration programs, including later perinecrotic proliferation (>24 h) (Bajt, Knight et al. 2003), CXCR2⁺ pro-proliferative responses (Nguyen, Umbaugh et al. 2022), and p21-mediated senescence (Umbaugh, Nguyen et al. 2024). Rather than acting as transitional cells, mid-zone hepatocytes deploy the Atf4–Chop–Btg2 axis to transiently redirect resources from cell division toward cytoprotective pathways, such as redox defense. This spatially restricted mechanism operates before overt necrosis expansion and, unlike earlier periportal/pericentral stress profiles (Umbaugh, Ramachandran et al. 2021), is causally supported by integrated spatial transcriptomics and functional validation (Cut&Run, AAV modulation). Our findings therefore bridge early stress adaptation with subsequent liver repair, revealing that mid-zone hepatocytes may preserve regenerative capacity by prioritizing survival over proliferation during the critical injury-initiation window.

The antiproliferative activity of Btg2 has been studied through an integrated network of transcriptional and post-transcriptional regulatory mechanisms. At the post-transcriptional level, Btg2 bridges poly(A)-binding protein cytoplasmic 1 (Pabpc1) with the CCR4-NOT deadenylase complex, accelerating degradation of cell cycle-related transcripts (Stupfler, Birck et al. 2016). Transcriptionally, Btg2 disrupts the cyclin B1–Foxm1 positive feedback loop, suppressing cyclin B1 expression and inducing G2/M arrest (Park, Kim et al. 2008). Btg2 also inhibits G1/S transition through cyclin-dependent kinase interactions and modulates p53 activity (Yuniati, Scheijen et al. 2019). In the context of APAP-induced liver injury, the stress-responsive Atf4–Chop pathway induces Btg2 expression in mid-zonal hepatocytes, where it likely coordinates a comprehensive proliferation arrest through these complementary mechanisms. This multilayered regulatory strategy enables hepatocytes to temporarily paused cell proliferation and redirect resources toward cellular repair and stress adaptation during the acute phase of toxic injury.

A key limitation of this study is the scarcity of early-phase APAP-induced liver injury human samples, which precludes definitive validation of the clinical relevance of these zonal adaptations. The apparent differences between the human and murine datasets are likely attributable to the stage of liver injury analyzed. Human samples were obtained from end-stage liver explants, in which regenerative programs are already active, whereas our mouse studies focused on the early (3–12 h) injury phase preceding overt regeneration. Consistent with this interpretation, one ST specimen displayed a stress-dominant mid-zonal transcriptional program resembling our murine observations, whereas the second exhibited a predominantly proliferative profile. These findings suggest that the balance between stress adaptation and hepatocyte proliferation is dynamic and strongly influenced by the temporal stage of liver injury. Although the available human datasets do not capture the early injury window examined in our mouse model, they support conservation of the mid-zonal stress-response program across species.

We also acknowledge several technical limitations of our spatial transcriptomics approach. While ST analysis and Ki67 staining consistently identified hepatocytes as the primary proliferating compartment during early APAP-induced liver injury, non-parenchymal cells (*e.g.*, immune cells, stellate cells) may contribute to division signals. Single-cell resolution techniques would be required to exclude these contributions fully. Additionally, our ST data alone do not independently confirm zonal proliferative restraint: ST lacks overt separation in S/G2/M scores, uses non-canonical markers (*e.g.*, *Nasp, Cks1b*), and has low sensitivity for rare Ki67⁺ events (∼1% of spots). Thus, our primary evidence for mid-lobular proliferation dynamics comes from Ki67 immunohistochemistry (Figure 1), with ST findings (Figure 2E) shown as exploratory and directionally consistent with transient impairment, but not as definitive proof of proliferative restraint.

In summary, our study uncovers a spatially coordinated stress-adaptation mechanism in mid-zone hepatocytes during early APAP-induced liver injury, where the Atf4–Chop axis transiently suppresses proliferation via Btg2 upregulation. By elucidating how hepatocytes prioritize survival before initiating repair, our work provides a framework for understanding the dynamic interplay between stress adaptation and regeneration in tissues.

## Materials and methods

### Animal experiments and procedures

*Animals C57BL/6J* mice (strain no. N000013) were used as wild-type (WT) mice. All mouse colonies were maintained at the Animal Core Facility of Yunnan University. The animal studies were approved by the Yunnan University Institutional Animal Care and Use Committee (IACUC, Approval No. YNU20220262). For APAP treatment, mice (8-12 weeks old) were fasted overnight (5:00pm to 9:00am) before *i.p.* injected with APAP (Sigma, A7085) at a dose of 300 mg/kg for male mice, as female mice are less susceptible to APAP-induced liver injury (Guerrero Munoz and Fearon 1984). Male mice have been the choice in the vast majority of the studies of APAP-induced liver injury reported in the literature (Campion, Johnson et al. 2008, Shan, Li et al. 2021, Ben-Moshe, Veg et al. 2022, Matchett, Wilson-Kanamori et al. 2024). Therefore, we used male mice in the majority of the experiments presented. In one experiment, mice were *i.p.* injected with various doses of APAP (100, 300, 500mg/kg). For carbon tetrachloride (CCl_4_) treatment, mice were *i.p.* injected with corn oil or CCl_4_ (Macklin, Cat #C805329) at a dose of 0.8ul/g (corn oil dilute to 20%).

In some experiments, APAP-treated mice were pre-injected intraperitoneally (*i.p.*) with either AAV-overexpression plasmid (AAV-TBG-Atf4/Ddit3/Btg2) or AAV-sgRNA knockdown plasmid (IP479-EFS-sgBtg2) for 3 weeks. AAV-TBG-EGFP and IP479-EFS-sgEGFP were used as controls, respectively. ALT measurement was performed using a diagnostic assay kit (Teco Diagnostics, Anaheim CA) to assess liver injury.

### Histology, Immunohistochemistry & immunofluorescence

#### H&E staining

Tissues were fixed overnight at 4°C using buffered 10% paraformaldehyde (Sangon Biotech, A500684-0500) and subsequently embedded in paraffin. For H&E staining, paraffin-embedded slides were deparaffinized and rehydrated following standard protocols. The sections were briefly immersed in hematoxylin for 30 seconds, rinsed in water, and then stained with Eosin solution for 3 minutes. After staining, the sections were dehydrated in ethanol, cleared in xylene, and mounted. Slides were examined under a microscope (Scan System, SQS-1000) and analyzed using ImageJ. Necrotic areas outlined by loss of cellular architecture on H&E.

#### Immunofluorescence on frozen section

Immunofluorescence staining was conducted on frozen sections of fresh liver tissues. Initially, the tissues were fixed using 2% paraformaldehyde for 1 h and subsequently dehydrated overnight in a 30% sucrose solution. The following day, tissue embedding was performed using OCT (SAKURA, 4583), after which ultra-thin sections of 5μm were sliced. Permeabilization was achieved using 0.2% Triton X-100 for 10 minutes at room temperature, followed by blocking with 5% normal goat serum (VivaCell, C2530-0100). Primary antibodies against mouse GS (Proteintech, 11037-2-AP, 1:200), Chop (Proteintech, 15204-1-AP, 1:1000), Cyp1a2 (Proteintech, 15204-1-AP, 1:50) and Ki67 (Abcam, ab180569, 1:1000) were then applied. Secondary antibodies (Affinipure Goat Anti-Rabbit 488-conjugated, Jackson ImmunoResearch, 111-545-003, 1:400; Goat anti-Rabbit IgG (H+L) Cross-Adsorbed Secondary Antibody, Alexa Fluor ™ 568, Invitrogen, A11011, 1:1000, Alexa Fluor 488-conjugated Affinipure Goat Anti-Mouse IgG+IgM(H+L), Jackson ImmunoResearch, 115-545-044, 1:400 and Alexa Fluor® 594 AffiniPure® Goat Anti-Mouse IgG (H+L), Jackson ImmunoResearch, 115-585-003, 1:200) were used accordingly. After washing, the slides were mounted using antifade medium, and nuclei were stained with DAPI (Beyotime, C1006, prediluted). Images were captured using an Olympus BP80 microscope and analyzed with ImageJ.

#### Immunohistochemistry

Livers were fixed in 10% buffered formalin for 24 hours before paraffin embedding. For immunohistochemical staining of paraffin-embedded liver tissue, 5 μm sections were prepared, deparaffinized, and subjected to antigen retrieval by microwaving for 20 minutes. Antigen retrieval was performed in pH 6.0 sodium citrate buffer (Ki-67, P62, Chop, Cyp1a2, Cyp2f2), pH 6.4 sodium citrate buffer (p-eIF2A), or pH 9.0 Tris-EDTA buffer (Atf4, Btg2). Sections were blocked with 3% hydrogen peroxide and permeabilized with 0.2% Triton X-100 for 10 minutes at room temperature, followed by blocking with 5% normal goat serum (VivaCell, C2530-0100) for 1 hour at room temperature. Primary antibodies were incubated overnight at 4°C at the following dilutions in blocking buffer: Ki-67 (Abcam, ab15580, 1:400), P62 (Abclonal, A19700, 1:1000), Cyp2f2 (SantaCruz, sc-374540, 1:200), Chop (Proteintech, 15204-1-AP, 1:200), Cyp1a2 (Santa Cruz, sc-53241, 1:50), p-eIF2A (CST, 3398S, 1:50), Atf4 (CST, 11815S, 1:200), and Btg2 (Proteintech, 22339-1-AP, 1:100). Secondary antibodies used were Goat anti-Rabbit IgG (H+L) Secondary Antibody, Biotin (Invitrogen, 65-6140, 1:1000) and peroxidase-AffiniPure Goat Anti-mouse IgG(H+L) (Jackson ImmunoResearch, 115-035-003, 1:1000). Avidin, NeutrAvidin™, Horseradish Peroxidase conjugate (Invitrogen, A2664, 1:2000) was used as the third antibody. After three washes with PBS, color development was achieved using the DAB Peroxidase Substrate Kit (ZSGB-BIO, ZLI-9018), followed by quenching in distilled water. Slides were counterstained with hematoxylin (BBI, A600701-0050), dehydrated to xylene, and images were captured using an Olympus BP80 microscope or Scan System SQS-1000 and analyzed with ImageJ.

### Western blot analysis

Mouse liver tissue (30 mg) was homogenized in 1 mL RIPA buffer (10 mM Tris–HCl pH 8.0, 1 mM EDTA, 0.5 mM EGTA, 1% Triton X-100, 0.1% sodium deoxycholate, 0.1% SDS, 140 mM NaCl, 1 mM PMSF, and protease inhibitor cocktail). The lysate was mixed with 5× SDS loading buffer (final 1×), boiled at 98°C for 10 min, and centrifuged at 12,000 rpm for 3 min at room temperature. Supernatants were separated by 10% SDS-PAGE, transferred to PVDF membranes, and blocked with 5% nonfat milk in TBST (Tris-buffered saline with 0.1% Tween-20). Membranes were incubated with primary antibodies against Cyp2e1 (Proteintech, 19937-1-AP, 1:5000) and GAPDH (Proteintech, 60004-1-Ig, 1:5000), followed by HRP-conjugated secondary antibodies: goat anti-mouse (Jackson ImmunoResearch, 115-035-003, 1:2000) and goat anti-rabbit (Jackson ImmunoResearch, 111-035-003, 1:2000). Protein bands were detected using enhanced chemiluminescence (ECL) and a Minichemi Chemiluminescence Imaging System.

### RNA extraction and quantitative real-time PCR

Ground mouse liver tissue was lysed in TRIzol reagent (Invitrogen, 15596018) and centrifuged at 12,000 rpm for 10 min at 4°C. The supernatant was transferred to a new tube, mixed with 200 μL chloroform, incubated at room temperature for 10 min, and centrifuged. The aqueous phase was collected, mixed with an equal volume of isopropanol, incubated at 4°C for 10 min, and centrifuged. The supernatant was discarded, and the RNA pellet was washed twice with 75% ethanol, air-dried for 10 min, and dissolved in RNase-free water. RNA concentration and purity were measured using a NanoDrop spectrophotometer. Reverse transcription was performed using a cDNA first-strand synthesis kit (Takara, 6210B). Quantitative PCR was carried out in triplicates using PowerUp™ SYBR™ Green Master Mix (Thermo Fisher, A25742) on a QuantStudio 1 Real-Time PCR system (Life Technologies) according to the manufacturer’s instructions. Primer sequences are provided in Table S1.

**Table S1.**
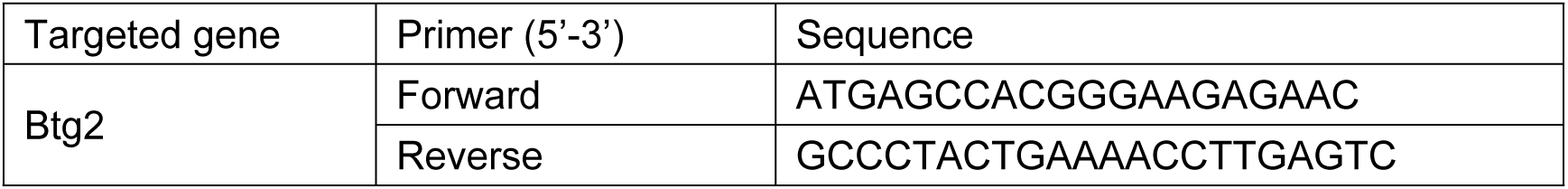
(qPCR primers)

### AAV production and purification

AAV8 was produced using 293T cells cultured in one or more 175 cm^2^ cell culture flasks. Cells were plated one day prior to transfection, reaching 50% confluence, which allowed them to reach 80-90% confluence the next day. For transfection of one 175 cm^2^ flask, a mixture of 11.56 μg pHelper, 7.24 μg pAAV2/8 (Addgene, 112864), and 5.9 μg transgene plasmids was prepared in 3 ml Opti-MEM medium in a tube. PEI solution (1 μg/μl in water, pH 7.0, powder from Polysciences, 23966-2) was added to the tube at a ratio of 3 μg PEI per 1 μg DNA. The solution was mixed and incubated for 15 minutes before adding to the cell culture. At 24 hours post-transfection, the medium was replaced with antibiotic-containing medium (P+/S+) without FBS. At 72 hours after the medium change, cells were scraped off the flask, and 3 ml of chloroform was added, followed by vortexing for 5 minutes. Then, 7.6 ml of 5M NaCl was added and vortexed for 10 seconds. The solution was centrifuged at 3000g for 5 minutes at 4°C. The supernatant was transferred to a new 50 ml tube, and 9.4 ml of 50% (w/v) PEG 8000 was added, vortexed for 10 seconds, and left for 1 hour at 4°C. Subsequently, the mixture was centrifuged at 3000g for 30 minutes at 4°C. The supernatant was decanted, and the tube was inverted for 10 minutes to dry. Next, 1.4 ml of 50 mM HEPES buffer (pH 8.0) was added to re-suspend the pellet, and the solution was vortexed for 5 minutes. Then, 3.5 μl of 1M MgCl_2_, 2.8 μl of 5 μg/μl DNase I, and 2.8 μl of 5 μg/μl RNase A were added, and the mixture was incubated for 20 minutes at 37°C. The virus was then extracted with chloroform, and the solvent was replaced with PBS. AAV was concentrated by centrifugation at 14000g for 5 minutes at 4°C using a 100kDa ultrafiltration tube (Beyotime, FUF058). Finally, the samples were aliquoted and stored at −80°C after titer measurement.

### Library preparation for 10x Visium spatial transcriptomics

After administering APAP, mice were euthanized at 0h, 3h, 6h and 24h time points using carbon dioxide. Freshly harvested liver tissues were frozen using isopentane and liquid nitrogen, embedded in OCT, sectioned into 10 μm slices, and mounted on Visium tissue optimization or spatial gene expression slides. Tissues were fixed with methanol, stained with H&E, and imaged at 40× magnification. Tissue permeabilization was optimized for 6 minutes. Libraries were prepared according to the Visium spatial gene expression user guide, with cDNA amplification cycles adjusted per time point, and index PCR performed for 6 cycles. Libraries were pooled, loaded at 400 pM, and sequenced on a NovaSeq 6000 SP100 sequencer.

### Processing of raw sequencing data

The raw data underwent processing using spaceranger (version 1.1.0) to perform several analytical tasks. These included detecting tissue boundaries, aligning reads to the reference genome, generating feature-spot matrices, conducting clustering to identify spatially distinct groups of cells, and analyzing gene expression patterns. Additionally, spaceranger facilitated the spatial placement of spots within the context of the slide image, providing a comprehensive spatial transcriptomic analysis of the tissue sample.

### Basic analysis of spatial transcriptomics (ST) data

For the gene-spot matrices generated by Spaceranger, we utilized Scanpy (version 1.9.3) for analysis (Patro, Duggal et al. 2017). Initially, routine statistical analyses were conducted, including calculating the number of detected UMIs (nUMI) and genes (nGene) in each spot. Basic quality control (QC) measures were then applied to the data. Specifically, spots with extremely low UMI or gene counts (min_counts=2000, min_cell=10) were excluded using scanpy.pp.filter_cells function.

Mitochondrial and hemoglobin genes were filtered out to focus on relevant gene expression. Spot matrix was filtered out to keep only spots overlaying tissue sections. Following QC, data normalization was performed using the scanpy.pp.normalize_total function, and log-transformation was applied using the scanpy.pp.log1p function. Subsequently, we identified 2000 highly variable genes using the scanpy.pp.highly_variable_genes function based on their expression means and variances.

Principal component analysis (PCA) was then applied to reduce the dimensionality of the data, projecting the spots into a lower-dimensional space using the identified principal components (PCs). Using the corrected PC matrices, we conducted unsupervised clustering based on shared nearest neighbors (SNN) and visualized the data using UMAP (Uniform Manifold Approximation and Projection) for further exploration and analysis (Lim and Qiu 2023).

Each sample was partitioned into three clusters. To facilitate comparison, each cluster was annotated with a region label (PC, PP, or Mid), determined by integrating information from cluster-specific marker genes and H&E staining images. Furthermore, to compare gene expression profiles across clusters, we identified differentially expressed genes using fold change analysis and the Wilcoxon rank sum test. This approach allowed us to highlight genes showing significant expression differences among all or selected clusters, providing insights into biological variations across spatial regions in the tissue samples.

### Analysis of differentially expressed genes (DEGs) and pathway enrichment

For differential gene analysis across different samples, we initially integrated all samples using the combat function to mitigate batch effects (Stuart, Butler et al. 2019). Subsequently, differential expression analysis of variable genes was conducted using the FindMarkers function, employing the Wilcoxon rank sum test (Ritchie, Phipson et al. 2015). Genes showing significant changes were selected based on criteria including a log-fold change threshold of 0.1 and a minimum percentage of samples showing differential expression of 1%. The differentially expressed genes (DEGs) identified were enriched across various time points post-APAP injection, relative to the 0-hour time point.

Pathway analysis was performed using the Kyoto Encyclopedia of Genes and Genomes (KEGG), while gene ontology (GO) analysis was carried out using the clusterProfiler package (Yu, Wang et al. 2012) to explore functional categories associated with these DEGs. The most significantly enriched pathways were identified using the enrichGO and enrichKEGG functions within the clusterProfiler package (Yu, Wang et al. 2012). Furthermore, Gene Set Variation Analysis (GSVA) was employed to investigate the expression variations related to proliferation at different time points following APAP administration, utilizing the gsva function from the GSVA R package (Hänzelmann, Castelo et al. 2013). Pathway activities were visualized using the pheatmap R package for comprehensive analysis and interpretation. To functionally classify the DEGs identified in the F2A comparison, we performed Gene Ontology (GO) enrichment analysis using clusterProfiler. Each DEG was mapped to GO terms from the three ontologies: biological process, cellular component, and molecular function. When a single gene was annotated with multiple GO terms, we selected the most prominent term—defined as the one with the smallest adjusted p-value or highest enrichment score—for downstream interpretation. This approach ensured that the primary functional role of each DEG was captured without redundancy.

### Gene regulatory network analysis

The Single-cell regulatory network inference and clustering (SCENIC) workflow comprises three main steps: coexpression analysis, target gene motif enrichment analysis, and regulon activity assessment. The analysis employed pySCENIC (version 0.12.1)(van de Sande, Flerin et al. 2020) with default parameters, using the raw count matrix from all samples as input. Co-expression modules were initially identified, and the interaction strength between transcription factors (TFs) and their target genes was evaluated using GRNBoost (Aibar, González-Blas et al. 2017). Each coexpressed module underwent motif enrichment analysis using RcisTarget with a rank threshold of 3000, to identify modules where the TF motif was significantly enriched among targets (Aibar, González-Blas et al. 2017). Only modules meeting these criteria were retained, establishing TFs along with their potential direct targets as regulons. The activity of each regulon within each cell was assessed using AUCell. For visualization, average regulon activity scores (AUC) were calculated for each cluster (Aibar, González-Blas et al. 2017). A rank plot of regulons was generated using ggplot2. Additionally, the gene expression network related to the target genes of the TFs of interest was visualized using the nx.draw function.

### Analysis of snRNA-seq Data (GSE223561)

To annotate cell clusters in the single-nucleus RNA-seq (snRNA-seq) data, we utilized a curated list of known marker genes corresponding to liver cell lineages, as reported in Multimodal Decoding of Human Liver Regeneration. Using the AddModuleScore function in Seurat, we calculated signature scores to assign cell lineage identities. Clusters identified as primarily composed of cycling cells were further reclustered to segregate them into their constituent lineages. This process was repeated iteratively for each identified lineage. A cleansing step was included to remove clusters enriched for nuclei identified as doublets or overexpressing marker genes from multiple lineages. Hepatocytes were subsequently extracted for further reclustering, using resolution (res) = 0.2 and 80 principal components (npc). To annotate the hepatocyte clusters, we applied a curated list of known zonation-specific marker genes for hepatocytes.

### Library preparation for Cut&Run samples

Cut&Run was conducted on mouse liver tissue samples from mice injected with APAP for 6 hours using the Cut&Run assay kit (CST, 86652) following the manufacturer’s protocol. Initially, 1 mg of fresh tissue was weighed for each antibody/MNase reaction, with an additional 1 mg for the input sample. Reactions included positive and negative controls. Tissues were fixed with 1 ml of 0.1% formaldehyde for 10 minutes and quenched with glycine for 5 minutes. After washing with PBS, tissues were resuspended in 1 ml of 1X Wash Buffer (+ spermidine + protease inhibitor cocktail) and transferred to a Dounce homogenizer. Tissue was homogenized into a single-cell suspension with 10-15 strokes until no tissue chunks remained.

For the Cut&Run assay to identify Chop binding sites, samples were bound to concanavalin A beads and incubated overnight at 4°C with the following primary antibodies: 0.5 μg CHOP (CST, 2895S, 1:100), 2 μl positive control Tri-Methyl-Histone H3 (Lys4) (CST, 9751), and 5 μl negative control Rabbit (DA1E) mAb IgG XP® Isotype Control (CST, 66362). The pAG MNase enzyme was activated by adding cold calcium chloride and incubating at 4°C for 30 minutes. The reaction was terminated with 1X stop buffer at 37°C for 10 minutes without shaking to release DNA fragments into the solution.

To reverse crosslinks in fixed tissue samples, samples were brought to room temperature and treated with 10% SDS Solution (BBI, B548118-0100) to a final concentration of 0.1%, along with proteinase K (20 mg/ml). DNA Extraction Buffer (+ Proteinase K + RNAse A) was added to the input sample. Cells were lysed and chromatin fragmented by sonication using a Covaris ME220 contact ultrasound apparatus on ice, with optimal conditions generating chromatin fragments ranging from 100 to 600 bp. DNA was purified using phenol/chloroform extraction followed by ethanol precipitation, and DNA quantification was performed by qPCR. For Cut&Run, purified DNA was used to prepare sequencing libraries with the NEB Next Ultra II DNA Library Prep Kit for Illumina (7645S and 7335L) and sequenced on an Illumina Nova-Seq 6000 sequencer to obtain 150 bp paired-end reads.

### Cut&Run data analysis

For visualization purposes, BW files were generated using Integrative Genomics Viewer (IGV) software (Robinson, Thorvaldsdóttir et al. 2011), and bigwig files were created using Deeptools (version 3.5.1) with the ‘bamCoverage’ module (bamCoverage --binSize 100) (Ramírez, Ryan et al. 2016). For each biological replicate and its corresponding IgG control, peaks were called using macs2 (version 2.2.7) with the command macs2 callpeak -f BAMPE --qvalue 0.1 --keep-dup (Zhang, Liu et al. 2008). To annotate the identified peaks, ChIPseeker (version 1.26.2) in R was utilized (Yu, Wang et al. 2015).

### Quantification of zonal distribution

The zonation of protein (such as Ki67, Chop, Atf4) -positive nuclei was quantified using the following method: the position index (P.I.) was calculated based on distances to the nearest CV (x), portal vein (PV) (y), and the distance between CV and PV (z), utilizing the law of cosines. The formula employed was P.I. = (x^2 + z^2 - y^2) / (2z^2). This approach aligns with the methodology described by Lin et al (Lin, Nascimento et al. 2018).

### Statistical analysis

All experimental data are presented as the mean ± SD. Statistical analyses were performed using GraphPad Prism (v8.0.2), with appropriate tests selected based on experimental design: unpaired two-tailed Student’s t-tests for comparisons between two groups, and one-way ANOVA for comparisons involving three or more groups. The number of animals (‘‘n’’) used in each experiment is indicated in the Figures and corresponding legends. For quantification of liver sections, three to five random pericentral and periportal fields of each liver sample, unless specified otherwise, were imaged and quantified using Image J.

## Supporting information

Key resources table

## Data and code availability

Cut&run data are accessible (GSE272565).

ST data are accessible (GSE272564).

This paper does not report original code.

## Acknowledgements

We thank Bin Qi (Yunnan University) for suggestions and discussion. We thank Dr. Yin Hao (Wuhan University) for providing us the pHelper and pAAV2/8 plasmids. We thank Hui Yang (Institute of Neuroscience, SBS, CAS) for providing us the plasmids related to CasRx system. We thank Yonglong Wei (Yunnan University) for helping in quantification of spatial distribution.

## Additional information

## Author Contributions

Y.Y.Z. performed the experiments, analyzed the data, and wrote the methods and figure legends. C.X.D. initially analyzed the sequencing data, while B.C. continued the sequencing data analysis, performed the experiments, wrote the methods and figure legends. J.H. prepared the ST samples. Y.N.L performed the experiments. S.L. performed some mice experiments. W.J.L. analyzed the sequencing data during revision. C.P. supervised the ST data analysis by C.X.D., participated in the initial study design, and revised the manuscript. Z.S. initiated, organized, and designed the study, and wrote the manuscript.

## Supplementary files

MDAR checklist

## Data availability

All data generated or analysed during this study are included in the manuscript and supporting files; source data files have been provided.

## Supplementary Figure

**Figure 2-figure supplement 1.**
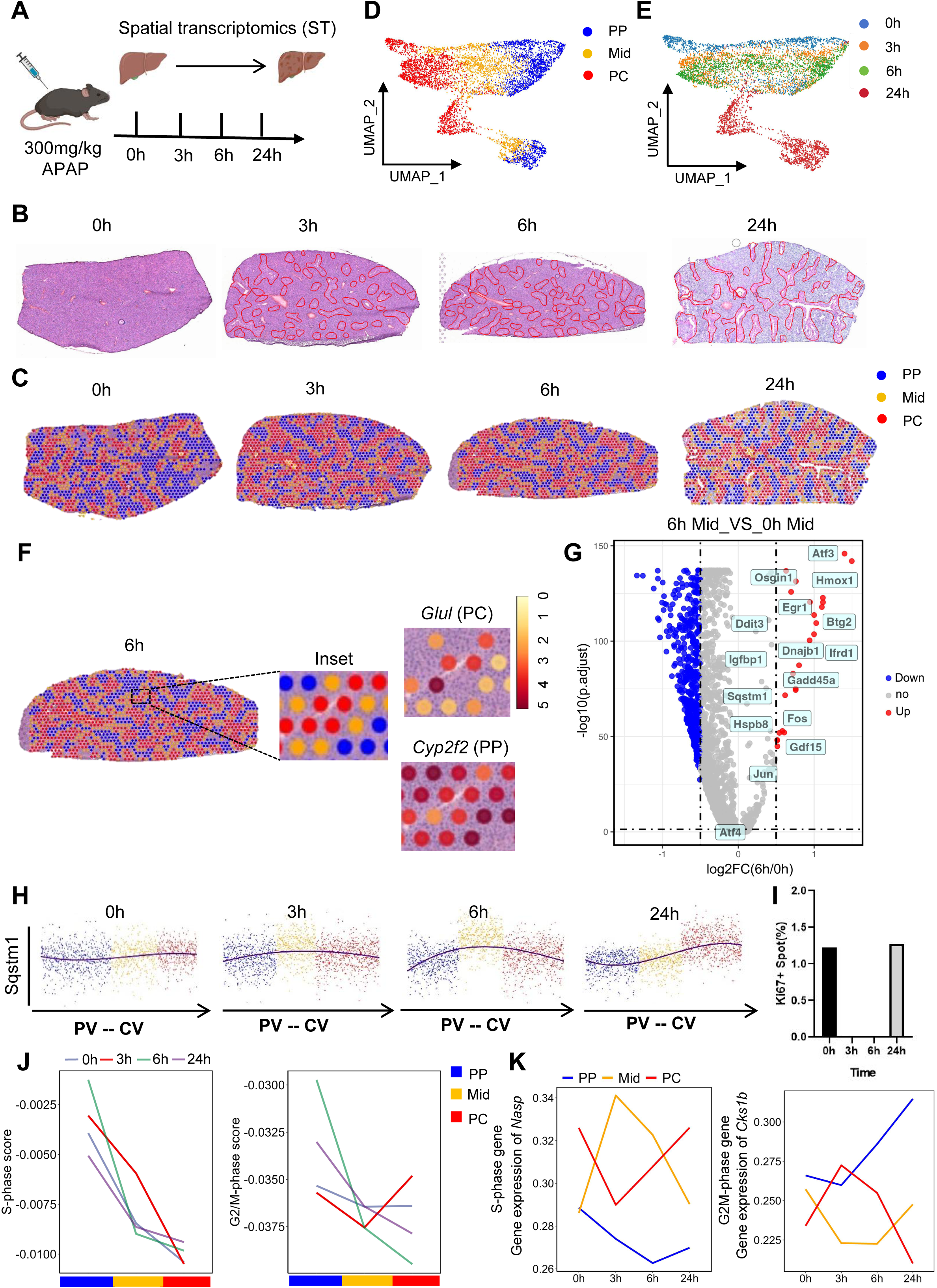
The liver displays transcriptome-wide zonation in acetaminophen-induced liver injury. **(A)** Schematic figure illustrating the experimental strategy for spatial transcriptome analysis. n=1 mouse/time point. **(B)** Representative H&E staining images illustrating liver morphology in mice following intraperitoneal injection of 300 mg/kg APAP at various time points (0, 3, 6 and 24 h). These liver sections were subsequently used for spatial transcriptome analysis. Injured area is outline by red line. Necrotic areas outlined by loss of cellular architecture on H&E. **(C)** Spatial map visualizing the spatiotemporal dynamics of hepatocyte zones (zones PP, Mid, and PC) at 0, 3, 6, 24 h post-APAP, respectively. **(D and E)** UMAP visualizing the spatiotemporal dynamics of hepatocyte zones (zones PP, Mid, and PC) at 0, 3, 6, 24 h post-APAP, based on zonal distribution and time points, respectively. **(F)** Spatiotemporally resolved heatmaps of representative PC marker Glul, PP marker Cyp2f2 in zoom-in area. **(G)** Volcano plot illustrating the DEGs in Mid-zone at 6 h compared to 0 h post-APAP. Gray dots denote genes that are not statistically significant. Red and blue dots represent genes that are upregulated and downregulated, respectively, in the sample tissue, at least a 0.5-fold difference from the matched control, with a false discovery rate (FDR) threshold of 0.05. **(H)** Scatter plot shows the dynamic changes of Sqstm1 mean expression log ₂ (TPM) in hepatocyte zones (zones PP, Mid, and PC) at 0 and 6 h post-APAP. TPM: Transcripts per million. **(I)** Percentage of Ki67-positive spots analyzed at 0, 3, 6 and 24 h post-APAP. **(J)** Average module score for S-phase (up) and G2/M-phase (down) genes across hepatocyte zones (zones PP, Mid, and PC) at 0, 3, 6, 24 h post-APAP. **(K)** Expression levels of the S-phase gene Nasp (up) and the G2/M phase gene Cks1b (down) in each hepatocyte zone (zones PP, Mid, and PC) at 0, 3, 6, 24 h post-APAP.

**Figure 3-figure supplement 1.**
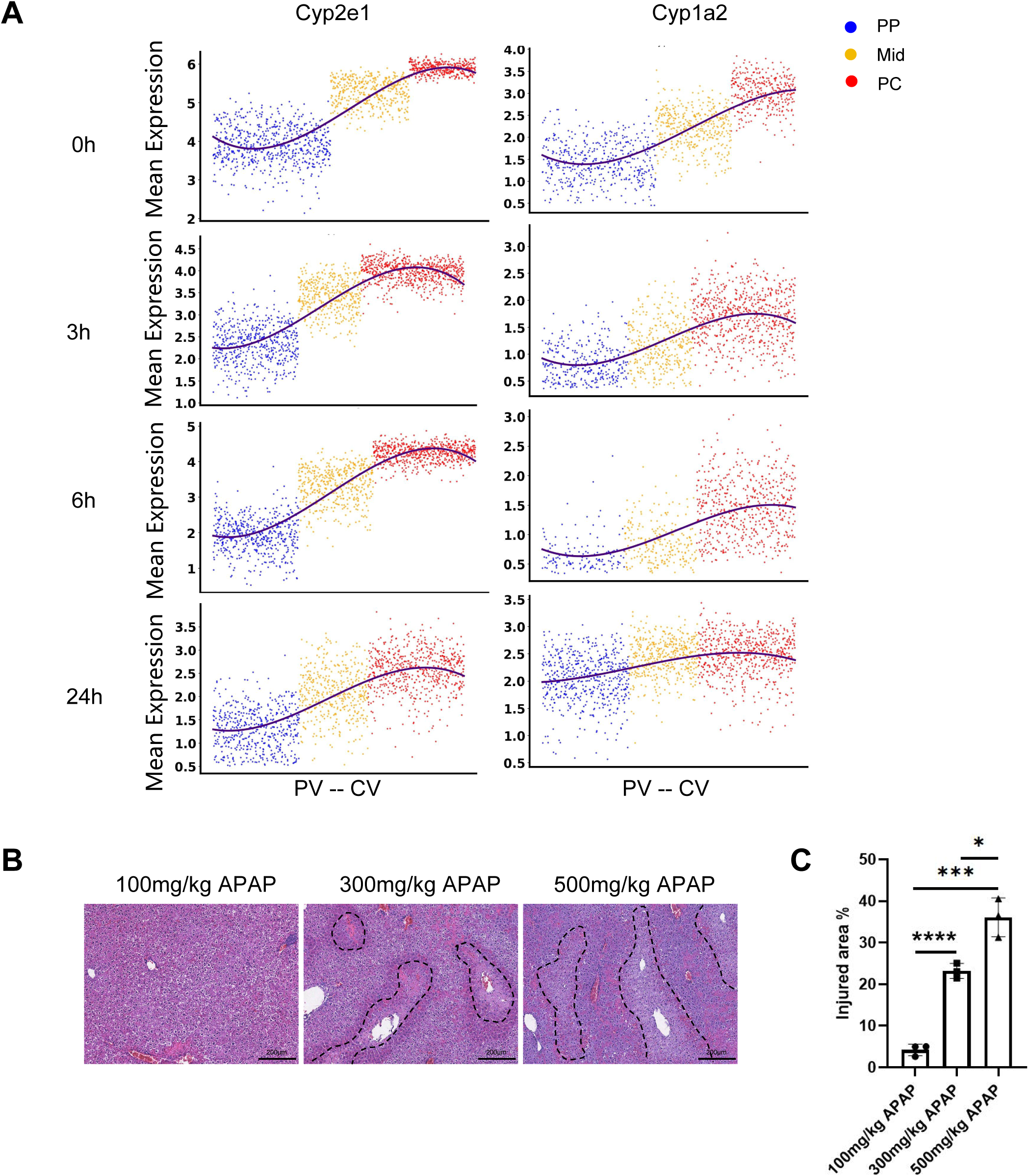
The zonal expression of the Cyp family dictates the zonal hepatocyte response to APAP. **(A)** Scatter plot showing the dynamic changes of Cyp2e1 and Cyp1a2 mean expression log ₂ (TPM), in hepatocyte zones (PP, Mid, PC) at 0, 3, 6, and 24 h post-APAP. TPM: Transcripts per million. **(B)** Representative H&E staining images illustrating liver morphology in mice following intraperitoneal injection of various doses of APAP (100, 300, 500mg/kg) at 6h post APAP. Injured area is outline by black dashed lines. Necrotic areas outlined by loss of cellular architecture on H&E. **(C)** The percentage of injured area is quantified. n=3 mice/group. Data are represented as means ± SD; One-way ANOVA (C). *p < 0.05; ***p < 0.001.

**Figure 4-figure supplement 1.**
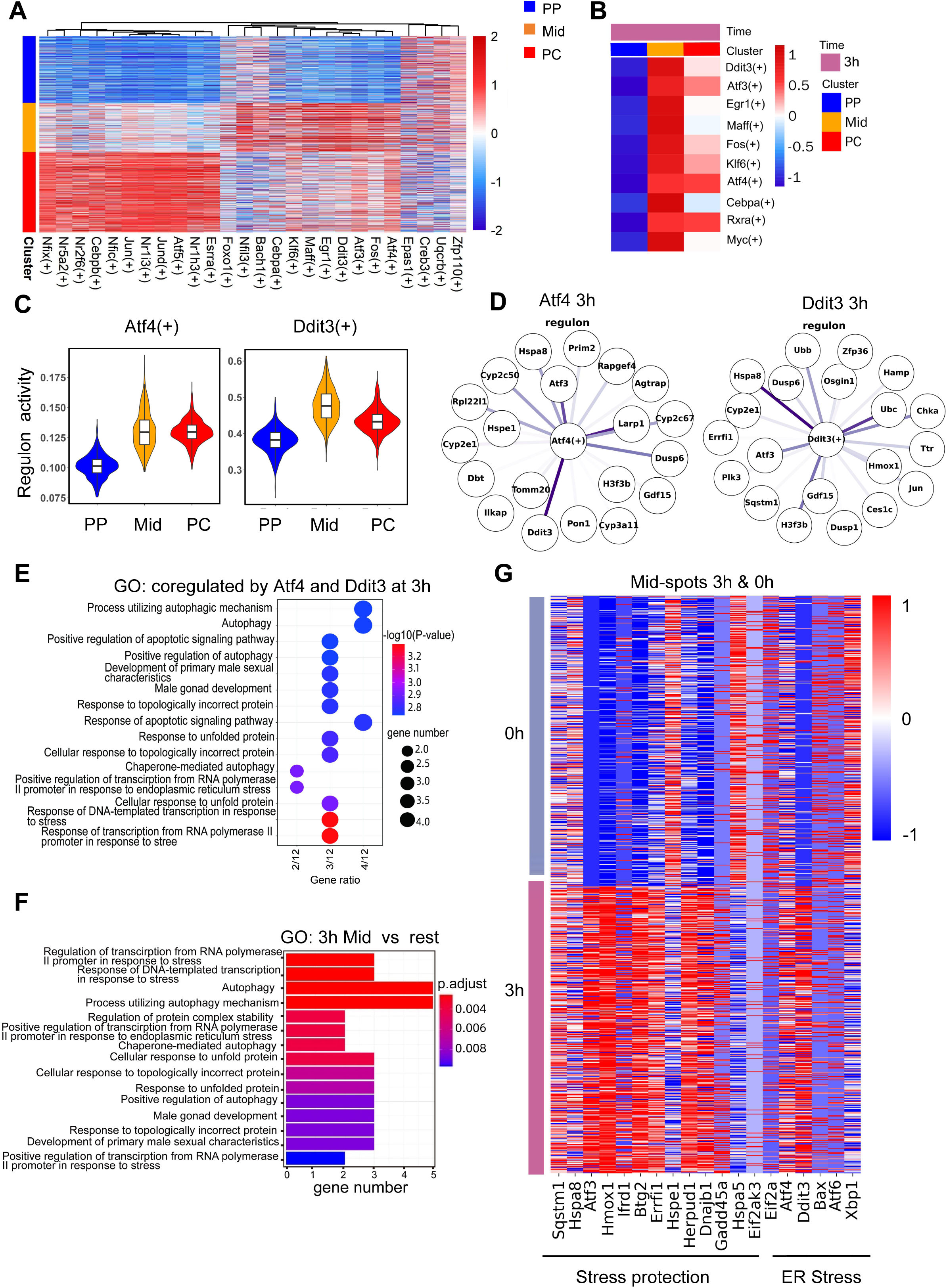
The Atf4-Ddit3 axis emerges as pivotal in mid-lobular hepatocytes during the initial stages of acute injury. **(A)** Heatmap displays the area under the curve (AUC) scores of transcription factor (TF) motifs, estimated per dot by Single-Cell Regulatory Network Inference and Clustering (SCENIC), highlighting gene regulatory networks and differentially activated TF motifs in hepatocyte zones (PP, Mid, PC) at 3 h post-APAP. Columns represent TF motifs, rows represent dots, and color intensity indicates AUC scores. **(B)** The heatmap shows inferred transcription factors (TFs) activity across different zonal regions at 3 h post-APAP. Activity was quantified as regulon enrichment scores using AUCell. The color scale represents regulon activity scores. **(C)** Violin plots shows regulon activity of Atf4 (left) and Ddit3 (right) in each zonation at 3 h post-APAP. Activity was quantified as regulon enrichment scores using AUCell. **(D)** Gene regulatory network displays Atf4 and its target genes, as well as Ddit3 and its target genes at 3 h post APAP, respectively. Edge color indicates the regulatory strength of transcription factors (TFs) on target genes, as measured by AUCell scores. **(E)** GO pathway analysis reveals the top 15 enriched pathways for genes co-regulated by Atf4 and Ddit3 in mid hepatocyte zones at 3 h post-APAP. **(F)** GO pathway analysis reveals the top 15 enriched pathways for up-regulated genes in Mid hepatocyte zones at 3 h compared to the rest hepatocyte zones post-APAP. **(G)** Heatmap displays stress response and ER stress-related genes from differentially expressed genes (DEGs) identified in Mid hepatocyte zones at 0 and 3 h post-APAP.

**Figure 4-figure supplement 2.**
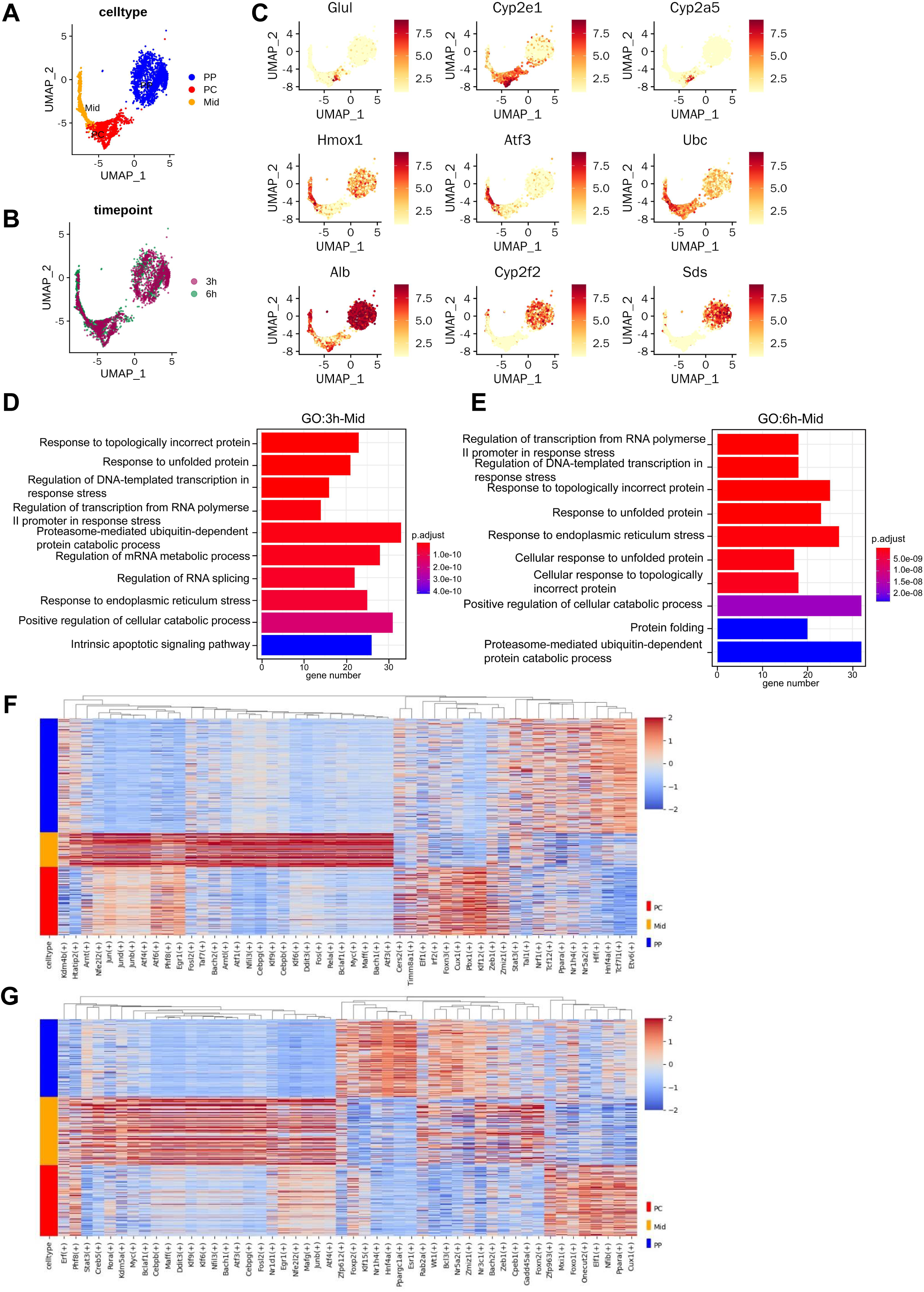
Validation of zonal hepatocyte responses and transcriptional changes post-APAP using the GSE223561 scRNA-seq dataset. **(A)** UMAP visualization of hepatocytes colored by cell type annotations. **(B)** UMAP visualization of hepatocytes colored by time post-APAP treatment. **(C)** UMAP visualization of hepatocytes colored based on the expression of centrally zonated genes (Glul, Cyp2e1, Cyp2a5), periportal genes (Alb, Cyp2f2, Sds), and mid-zonal genes (Hmox1, Atf3, Ubc). **(D)** GO pathway analysis showing the top 10 enriched pathways for upregulated genes in mid-zonal hepatocytes at 3 h post-APAP. **(E)** GO pathway analysis showing the top 10 enriched pathways for upregulated genes in mid-zonal hepatocytes at 6 h post-APAP. **(F)** Heatmap illustrating the area under the curve (AUC) scores of transcription factor (TF) motifs (Top 50), as estimated per cell using Single-Cell Regulatory Network Inference and Clustering (SCENIC). Differentially activated motifs in each zonal region are shown for 3 h post-APAP. **(G)** Heatmap illustrating the AUC scores of TF motifs (Top 50) estimated per cell using SCENIC, highlighting differentially activated motifs in each zonal region at 6 h post-APAP.

**Figure 4-figure supplement 3.**
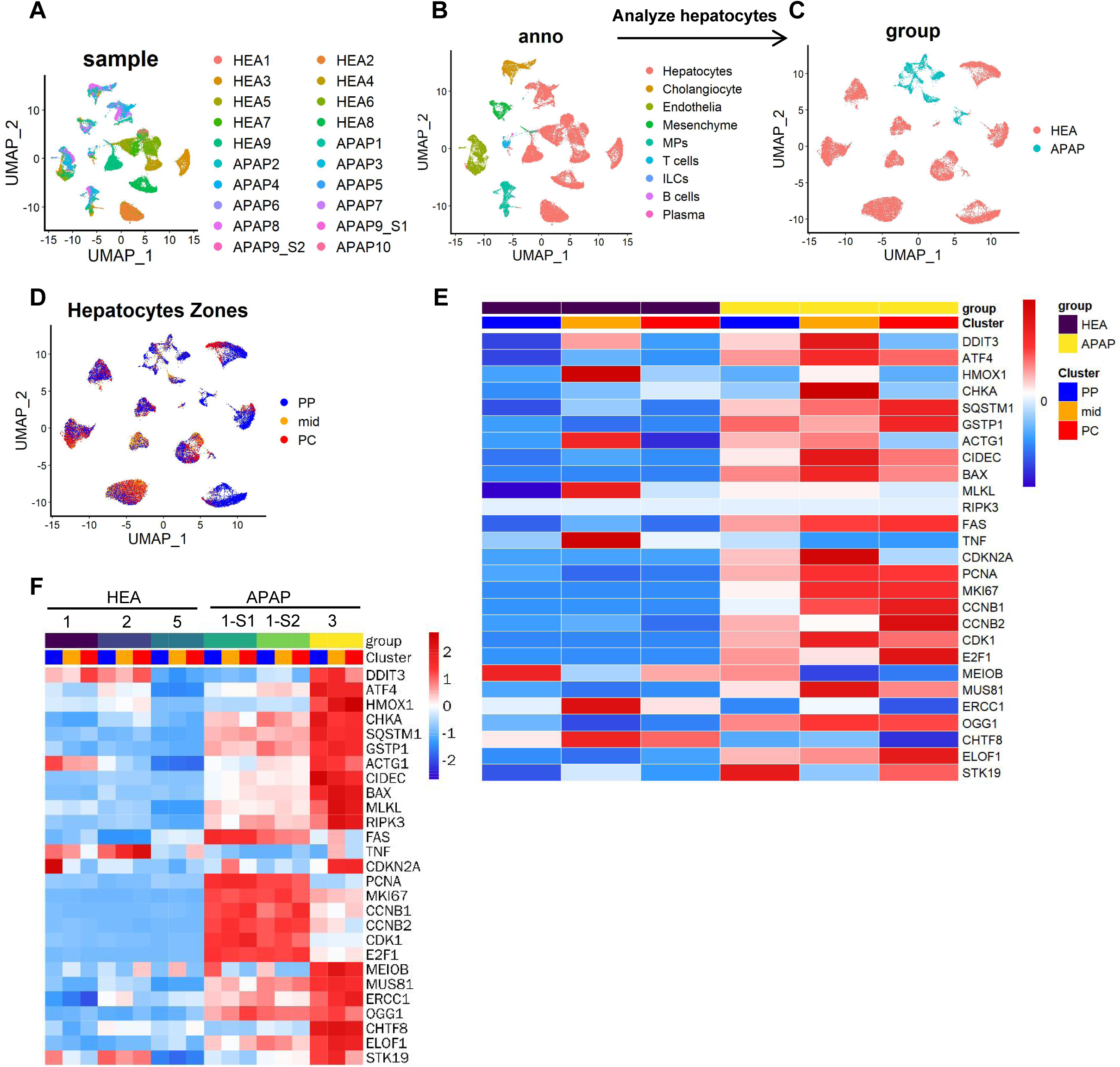
Zonal hepatocyte responses and transcriptional reprogramming in human APAP-induced liver injury, analyzed using the GSE223561 dataset. **(A)** UMAP projection of all single-nucleus RNA-sequencing (snRNA-seq) nuclei, colored by individuals (healthy controls versus APAP patients; n=9 and n=10, respectively). **(B)** UMAP projection of all snRNA-seq nuclei, annotated by major hepatic cell lineages. **(C)** UMAP projection of the hepatocyte subset, colored by disease status. **(D)** UMAP projection of hepatocytes showing expression levels of representative zonation markers: pericentral (Glul, Cyp2e1, Cyp2a5), periportal (Alb, Cyp2f2, Sds), and mid-zonal (Igfbp2, Hamp, Hamp2). **(E)** Heatmap comparing zonal expression patterns of genes involved in stress response/cell death, cell cycle/proliferation, and DNA repair across periportal (PP), mid-zonal (Mid), and pericentral (PC) hepatocyte clusters in healthy and APAP groups from the snRNA-seq dataset. **(F)** Heatmap showing the spatial distribution of the same gene categories across PP, Mid, and PC zones in healthy and APAP patient samples from the spatial transcriptomics (ST) dataset.

**Figure 6-figure supplement 1.**
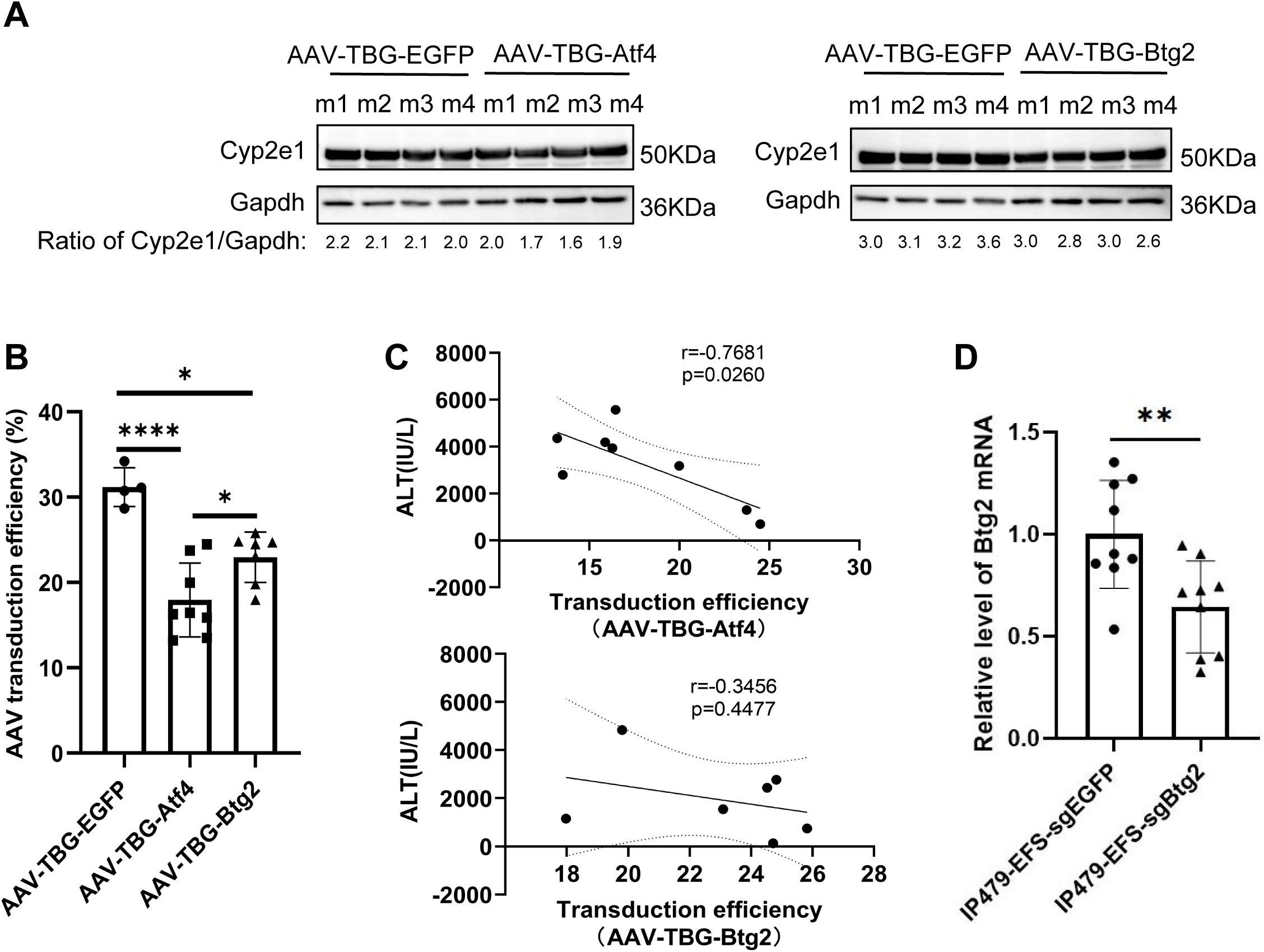
Transduction efficiency and confounding effects of AAV-TBG overexpression. **(A)** Western blot analysis of Cyp2e1 protein in whole liver lysates from mice transduced with 1.2 × 10¹¹ viral genome copies of AAV-TBG-EGFP, AAV-TBG-Atf4, or AAV-TBG-Btg2. Cyp2e1/Gapdh ratios are shown. n = 4 mice/group. **(B)** Transduction efficiency of AAV-TBG-EGFP, AAV-TBG-Atf4, and AAV-TBG-Btg2 in mouse liver was measured by immunostaining for the transgene proteins. n = 4–8 mice/group. **(C)** Correlation between transduction efficiency of AAV-TBG-Atf4 or AAV-TBG-Btg2 and serum ALT levels in the respective AAV-injected mice. Pearson correlation coefficient r and p values are indicated. **(D)** Btg2 mRNA expression in liver tissues from IP479-EFS-sgEGFP and IP479-EFS-sgBtg2 mice. n=9 mice/group. Data are represented as means ± SD; Pearson correlation (C); Unpaired two tailed Student’s t-test (B, D). *p < 0.05; **p < 0.01; ****p < 0.0001.

**Figure 6-figure supplement 2.**
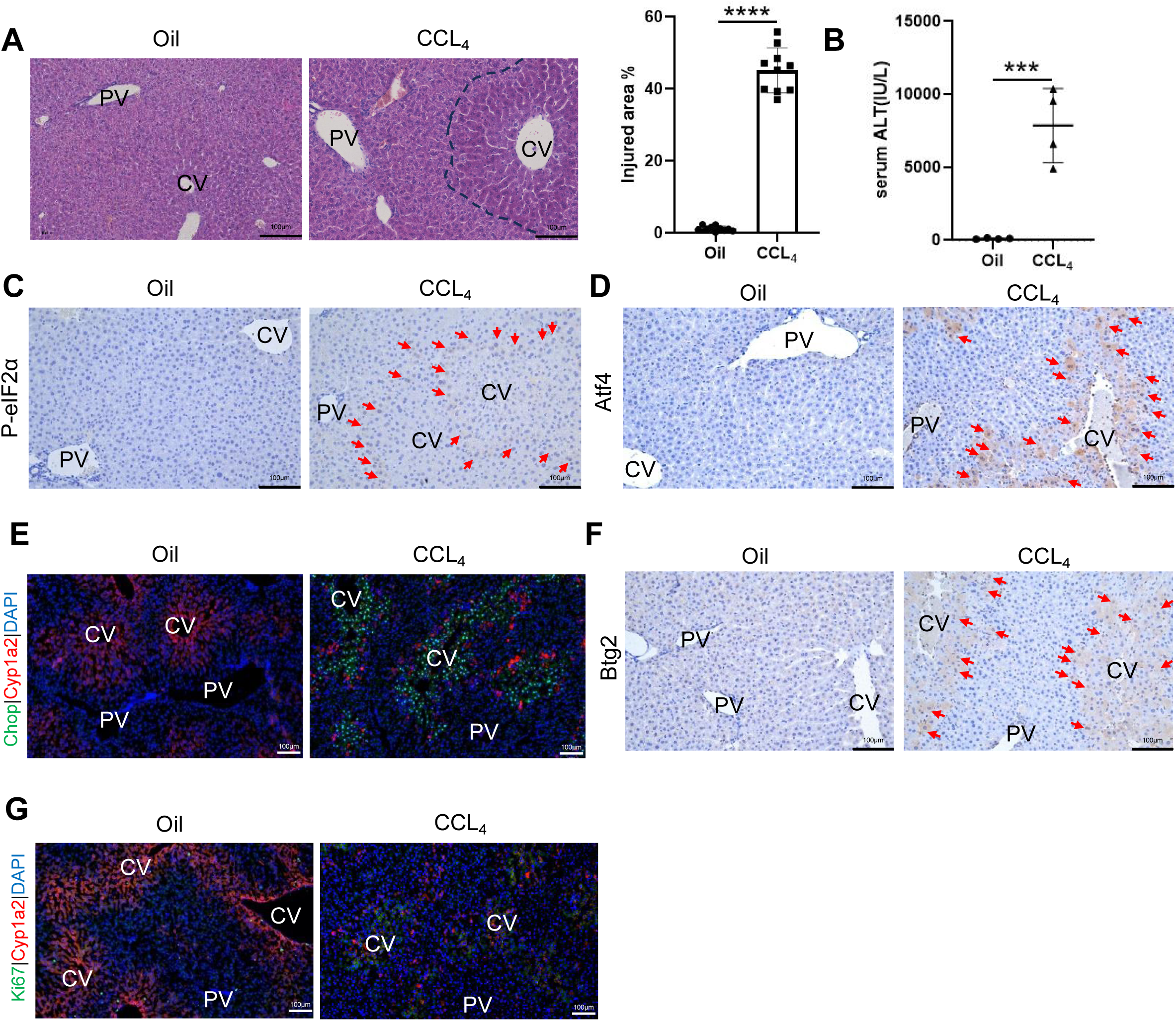
The ISR-Btg2 axis emerges in CCL4-induced acute liver injury. **(A)** H&E staining shows morphology of livers from mice at 18 h post oil and CCL4 injection. Injured area is outlined by black dashed lines. The percentage of injury area is quantified. n=4 mice/group. Necrotic areas outlined by loss of cellular architecture on H&E. **(B)** Serum levels of ALT is measured at 18h post-CCL4 mice. n=4 mice/group. **(C)** Immunohistochemistry staining of p-eIF2A in liver sections from mice at 18 h post oil and CCL4 injection. p-eIF2A-positive hepatocytes are indicated by red arrows. n=4 mice/group. **(D)** Immunohistochemistry staining of Atf4 in liver sections from mice at 18 h post oil and CCL4 injection. Atf4 -positive hepatocytes are indicated by red arrows. n=4 mice/group. **(E)** Immunofluorescence staining of Chop protein (green) in liver sections from mice at 18 h post oil and CCL4 injection. Cyp1a2 protein (red) staining highlights the area around the CV. Cell nuclei are stained with DAPI (blue). n=4 mice/group. **(F)** Immunohistochemistry staining of Btg2 in liver sections from mice at 18 h post oil and CCL4 injection. Btg2 -positive hepatocytes are indicated by red arrows. n=4 mice/group. **(G)** Immunofluorescence staining of Ki67 (red) was performed to assess proliferating hepatocytes in mice at 18 h post oil and CCL4 injection. Cyp1a2 protein (red) staining highlights the area around the CV. Cell nuclei were stained with DAPI (blue). n=4 mice/group. Data are represented as means ± SD; Unpaired Student’s t-test (A, B). *p < 0.05; **p < 0.01; ***p < 0.001; ****p < 0.0001; ns, not significant.

