## Supplementary material for "Mid-zone hepatocytes trade proliferation for survival via Atf4-Chop axis in early acute liver injury": Key resources table

| **Key resources table** | | | | |
| --- | --- | --- | --- | --- |
| **Reagent type (species) or resource** | **Designation** | **Source or reference** | **Identifiers** | **Additional information** |
| Antibody | Rabbit polyclonal anti-Ki67 antibody | Abcam | Cat# ab15580; RRID: N/A | IF(1:1000)  IHC(1:400) |
| Antibody | Rabbit Glutamine Synthetase Polyclonal antibody | Proteintech | Cat# 11037-2-AP; RRID: AB_2110650 | IF(1:200) |
| Antibody | Mouse monoclonal Anti-Cyp1a2 antibody | SantaCruz | Cat# sc-53241; RRID: N/A | IF(1:50)  IHC(1:50) |
| Antibody | Rabbi CHOP; GADD153 Polyclonal antibody | Proteintech | Cat# 15204-1-AP; RRID: AB_2292610 | IF(1:1000)  IHC(1:200) |
| Antibody | Monoclonal ATF-4 (D4B8) Rabbit mAb | Cell Signalling Technology | Cat# 11815S; RRID: N/A | IHC(1:200) |
| Antibody | Rabbit BTG2 Polyclonal antibody | Proteintech | Cat# 22339-1-AP; RRID: AB_2879078 | IHC(1:100) |
| Antibody | Polyclonal Phospho-eIF2α (Ser51) (D9G8) XP Rabbit mAb | Cell Signalling Technology | Cat# 3398s; RRID: N/A | IHC(1:50) |
| Antibody | Goat anti-Rabbit IgG (H+L) Secondary Antibody, Biotin | Invitrogen | Cat# 65-6140; RRID: AB_2533969 | IHC(1:1000) |
| Antibody | Peroxidase-conjugated Affinipure Goat Anti-Mouse IgG(H+L) | Jackson ImmunoResearch | Cat# 115-035-003; RRID: AB_10015289 | IHC(1:1000)  WB(1:2000) |
| Antibody | Peroxidase-conjugated Affinipure Goat Anti-Rabbit IgG(H+L) | Jackson ImmunoResearch | Cat# 111-035-003; RRID: AB_2313567 | WB (1:2000) |
| Antibody | Avidin, NeutrAvidin™, Horseradish Peroxidase conjugate | Invitrogen | Cat# A2664; RRID: N/A | IHC(1:2000) |
| Antibody | Alexa Fluor 488-conjugated Affinipure Goat Anti-Mouse IgG+IgM(H+L) | Jackson ImmunoResearch | Cat# 115-545-044; RRID: AB_2338844 | IHC(1:400) |
| Antibody | Alexa Fluor® 594 AffiniPure® Goat Anti-Mouse IgG (H+L) | Jackson ImmunoResearch | Cat# 115-585-003; RRID: AB_2338871 | IHC(1:200) |
| Antibody | Alexa fluor 568-goat anti-rabbit IgG(H+L)cross-adsorbed secondary antibody | Invitrogen | Cat# A11011; RRID: AB_143157 | IHC(1:1000) |
| Antibody | Alexa Fluor 488 AffiniPure™ Goat Anti-Rabbit IgG (H+L) | Jackson ImmunoResearch | Cat# 111-545-003; RRID: AB_2338046 | IHC(1:400) |
| Antibody | CYP2E1-Specific Polyclonal antibody | Proteintech | Cat# 19937-1-AP; RRID: AB_10646444 | WB(1:5000) |
| Antibody | GAPDH Monoclonal antibody | Proteintech | Cat# 60004-1-Ig; RRID: AB_2107436 | WB(1:5000) |
| Chemical compound, drug | Acetaminophen(APAP) | Sigma | Cat# A7085 |  |
| Chemical compound, drug | Paraformaldehyde, (CH2O)n | Sangon Biotech | Cat# A500684-0500 |  |
| Chemical compound, drug | Phosphate Buffered Saline | VivaCell | Cat# C3580-0500 |  |
| Chemical compound, drug | 10% Neutral Formalin Fix Solution | Sangon Biotech | Cat# E672001-0500 |  |
| Chemical compound, drug | Sucrose, C12H22O11 | Sangon Biotech | Cat# A502792-0005 |  |
| Chemical compound, drug | High effect paraffin cere sin | Shanghai Hualing Rehabilitation Equipment Manufacturing Plant. | Cat# N/A |  |
| Chemical compound, drug | Xylene | Tianjin Zhiyuan Chemical Reagents Co., Ltd. | Cat# N/A |  |
| Chemical compound, drug | Ethanol | Tianjin Zhiyuan Chemical Reagents Co., Ltd. | Cat# N/A |  |
| Chemical compound, drug | Citrate Antigen Retrieval Solution (Powder) | Sangon Biotech | Cat# E673002-0020 |  |
| Chemical compound, drug | Tris | Solarbio | Cat# T8060 |  |
| Chemical compound, drug | Disodium salt dihydrate, C10H14N2O8Na2·2H2O(EDTA) | Sangon Biotech | Cat# A500838-0500 |  |
| Chemical compound, drug | Hydrogen peroxide |  |  |  |
| Chemical compound, drug | Triton X-100 | Sangon Biotech | Cat# A600198-0500 |  |
| Chemical compound, drug | Goat serum | VivaCell | Cat# C2530-0100 |  |
| Chemical compound, drug | Hematoxylin | Sangon Biotech | Cat# A600701-0050 |  |
| Chemical compound, drug | Eosin Y(water soluble) | [Aladdin](https://www.aladdin-e.com/" \t "_blank) | Cat# E141405 |  |
| Chemical compound, drug | Neutral balsam | Solarbio | Cat# G8590 |  |
| Chemical compound, drug | Tissue-tek OCT compound | SAKURA | Cat# REF:4583 |  |
| Chemical compound, drug | Acetone | Chron Chemicals | Cat# N/A |  |
| Chemical compound, drug | Tween20 | Sangon Biotech | Cat# A600560-0500 |  |
| Chemical compound, drug | DAPI Staining Solution | Beyotime | Cat# C1006 |  |
| Chemical compound, drug | Isopentane | [Aladdin](https://www.aladdin-e.com/" \t "_blank) | Cat# M108171 |  |
| Chemical compound, drug | Methanol | Tianjin Zhiyuan Chemical Reagents Co., Ltd. | Cat# N/A |  |
| Chemical compound, drug | 20X TBS buffer | Sangon Biotech | Cat# B548105-0500 |  |
| Chemical compound, drug | UltraPure™ DNase/RNase-Free Distilled Water | Invitrogen | Cat# 10977015 |  |
| Chemical compound, drug | Trichloromethane | Chron Chemicals | Cat# N/A |  |
| Chemical compound, drug | FBS | VivaCell | Cat# C04001-500 |  |
| Chemical compound, drug | DMEM(High glucose) | VivaCell | Cat# C3113-0500 |  |
| Chemical compound, drug | Penicillin-Streptomycin Solution | VivaCell | Cat# C3421-0100 |  |
| Chemical compound, drug | Trypsin 1:300 from Porcine pancreas | Sangon Biotech | Cat# A100260-0050 |  |
| Chemical compound, drug | PEI | Polysciences | Cat# 23966-2 |  |
| Chemical compound, drug | Opti-MEM | Gibco | Cat# 11058021 |  |
| Chemical compound, drug | PEG-8000 | Sangon Biotech | Cat# A600433-0500 |  |
| Chemical compound, drug | MgCl2 | Ghtech | Cat# N/A |  |
| Chemical compound, drug | 1M HEPES | Solarbio | Cat# H1095 |  |
| Chemical compound, drug | 37% formaldehyde | Sigma | Cat# F8775 |  |
| Chemical compound, drug | 10% SDS Solution | Sangon Biotech | Cat# B548118-0100 |  |
| Chemical compound, drug | Glycine | Sangon Biotech | Cat# A502065-0005 |  |
| Chemical compound, drug | Pierce™ Protease inhibitor tablets, EDTA free | Thermo | Cat# A32965 |  |
| Chemical compound, drug | DNA extract buffer | Solarbio | Cat# P1012 |  |
| Chemical compound, drug | 3M Sodium acetate | Invitrogen | Cat# AM9740 |  |
| Chemical compound, drug | Glycogen | Thermo | Cat# R0561 |  |
| Chemical compound, drug | SSC buffer 20X concentrate | Sigma | Cat# S6639 |  |
| Chemical compound, drug | PowerUp™ SYBR™ Green Master Mix | Applied biosystems | Cat# A25742 |  |
| Chemical compound, drug | Proteinase K Solution (20 mg/ml) | Sangon Biotech | Cat# B600169-0002 |  |
| Chemical compound, drug | DNaseⅠ ,RNase-free | Thermo | Cat# EN0521 |  |
| Chemical compound, drug | Rnase A | Thermo | Cat# R1253 |  |
| Chemical compound, drug | Agencourt® AMPure® XP magnetic beads | Beckman Coulter | Cat# A63880 |  |
| Chemical compound, drug | Ethyl alcohol, Pure (200 proof, molecular biologygrade) | Sigma-Aldrich | Cat# E7023-500ML |  |
| Chemical compound, drug | DEPC-treated water | Biosharp | Cat# 701062 |  |
|  | **Critical commercial assays** | | |  |
| Critical commercial assays | Visium Spatial Gene Expression Slide & Reagent Kit, 4 rxns | 10X Genomics | Cat# PN-1000187 |  |
| Critical commercial assays | Visium Accessory Kit | 10X Genomics | Cat# PN-1000194 |  |
| Critical commercial assays | Dual Index TT Set APN | 10X Genomics | Cat# PN-1000215 |  |
| Critical commercial assays | Alanine aminotransferase assay kit | Nanjing Jiancheng Bioengineering Institute | Cat# c009－2－1 |  |
| Critical commercial assays | Aspartate aminotransferase Assay Kit | Nanjing Jiancheng Bioengineering Institute | Cat# C010-2-1 |  |
| Critical commercial assays | CUT&RUN Assay Kit | Cell Signalling Technology | Cat# 86652 |  |
| Critical commercial assays | NEBNext Ultra II DNA Library Prep Kit for Illumina | New England Biolabs | Cat# 7645S |  |
| Critical commercial assays | NEBNext Multiplex Oligos for Illumina (Index Primers Set 1) | New England Biolabs | Cat# 7335L |  |
| Critical commercial assays | Qubit™ dsDNA HS Assay Kit | Thermo Fisher  Scientific | Cat# Q32851 |  |
| Critical commercial assays | 2 × Taq Master Mix (Dye Plus) | Vazyme | Cat# P112-01 |  |
| Critical commercial assays | 2 × Rapid Taq Master Mix | Vazyme | Cat# P222-01 |  |
| Cell line | 293T | This paper | N/A |  |
| Deposited data | Spatial transcriptomics of mouse liver cells at different time points after APAP injection | This paper | GSE272564 |  |
| Deposited data | Cut&Run data of APAP-injured hepatocytes | This paper | GSE272565 |  |
| Strain, strain background (Mus musculus, male) | C57BL/6J | This paper | Strain ID N000013 |  |
| Software and algorithms | GraphPad Prism | GraphPad Software | https://www.graphpad.com |  |
| Software and algorithms | SPSS |  |  |  |
| Software and algorithms | ImageJ | National Institutes of Health | https://imagej.nih.gov/ij/ |  |
| Software and algorithms | Space ranger (v1.2.2) | 10x Genomics | https://10xgenomics.com |  |
| Software and algorithms | ClusterProfiler R package (v4.1.4) | Yu et al. | https://bioconductor.org/packages/release/bioc/html/clusterProfiler.html |  |
| Software and algorithms | Seurat R package (v4.2.3) | Stuart et al. | https://satijalab.org/seurat/ |  |
| Software and algorithms | Bowtie2 (v2.2.5) | Langmead et al. | <http://bowtie-bio.sourceforge.net/bowtie2/index.shtml> |  |
| Software and algorithms | Deeptools (v3.5.1) | Ramirez et al. | <https://deeptools.readthedocs.io/> |  |
| Software and algorithms | Integrative Genomics Viewer (IGV) | Robinson et al. | <https://igv.org/> |  |
| Software and algorithms | R (v4.0.5)and R studio | R Consortium | <https://www.rstudio.com/> |  |
| Software and algorithms | Python(v3.7.1) | Python | <https://www.python.org/> |  |
| Software and algorithms | GSVA R package (v1.38.2) | Hanzelmann et al. | https://www.bioconductor.org/packages/release/bioc/html/GSVA.html |  |
| Software and algorithms | Limma R package (v3.46.0) | Ritchie et al. | https://www.bioconductor.org/packages/release/bioc/html/limma.html |  |
| Software and algorithms | Pheatmap R package (v.1.0.12) | N/A | https://ggplot2.tidyverse.org |  |
| Software and algorithms | Ggplot2 R package | N/A | https://ggplot2.tidyverse.org |  |
| Software and algorithms | Scanpy (python package) | Wolf et al. | https://scanpy.readthedocs.io |  |
| Software and algorithms | pySCENIC (version 0.12.1) | Van de Sande B et al. | https://scenic.aertslab.org |  |
| Software and algorithms | Stellaris FISH Probe Designer | Biosearch Technology | www.biosearchtech.com |  |
